# Endogenous and fluorescent sterols reveal the molecular basis for ligand selectivity of human sterol transporters

**DOI:** 10.1101/2024.07.22.604041

**Authors:** Laura Depta, Hogan P. Bryce-Rogers, Nienke J. Dekker, Anna Wiehl Bønke, Nicolo’ Camporese, Mingxing Qian, Yuanjian Xu, Douglas F. Covey, Luca Laraia

## Abstract

Sterol transport proteins (STPs) play a pivotal role in cholesterol homeostasis and therefore are essential for healthy human physiology. Despite recent advances in dissecting functions of STPs in the human cell, there is still a significant knowledge gap regarding their specific biological functions and a lack of suitable selective probes for their study. Here, we profile fluorescent steroid-based probes across ten STPs, uncovering substantial differences in their selectivity, aiding the retrospective and prospective interpretation of biological results generated with those probes. These results guided the establishment of an STP screening panel combining diverse biophysical assays, enabling the evaluation of 41 steroid-based natural products and derivatives. Combining this with a thorough structural analysis revealed the molecular basis for STP specific selectivity profiles, leading to the uncovering of several new potent and selective Aster-B inhibitors, and supporting the role of this protein in steroidogenesis.

## INTRODUCTION

The regulation of intracellular cholesterol homeostasis is essential for a healthy human physiology. The functions of this important lipid are diverse, from controlling membrane structure and fluidity to serving as precursors to steroid hormones.^1^ Intracellular sterol transport proteins (STPs) are responsible for the non-vesicular transport of cholesterol and other sterols between specific organelles, which are divided into three protein families: the ORPs, the STARDs and the Asters.^2–4^

The oxysterol binding protein (OSBP) and OSBP-related proteins (ORPs) are key mediators and regulators of lipid transport between the endoplasmic reticulum (ER) and other organelles.^5,6^ They share a conserved OSBP-related domain (ORD), which has been shown to bind and transfer sterols and other lipids including phosphatidylcholine (PC), ceramide (CE) and phosphatidylinositol phosphates (PIPs).^7^ The mammalian Steroidogenic Acute Regulatory Protein (StAR)-related Lipid Transfer (START) Domain (STARD) family contains 15 members. All members have the StART lipid transfer domain, while only the membrane targeted STARD1/D3 subfamily and the soluble STARD4 subfamily (STARD4/D5/D6) are reported to bind sterols.^8^ While STARD1 regulates the delivery of cholesterol from the outer (OMM) to the inner mitochondrial membrane (IMM) in steroidogenesis, STARD3 transfers cholesterol from the ER to the late endosomes and mediates interactions between those two organelles.^9^ The Asters transport cholesterol from the PM to the ER, however, different expression patterns suggest specific functions.^10,11^ Aster-A is highly expressed in the brain, Aster-B in adrenal tissues, and Aster-C in the testis and liver. Aster-A and -C both play a role in autophagosome biogenesis, while Aster-B has been shown to regulate mitochondrial sterol transport and Aster-C has been shown to regulate mTOR activity.^12–14^

So far, information about the specific functions and transport mechanisms within the STP families is mainly based on knock-down and knock-out studies, which sometimes show contradictory results due to their documented functional redundancy.^15^ Here, an investigation with chemical tools would help to further elucidate the complex interplay of this network. Such a strategy necessitates bioactive ligands with a defined selectivity profile for each protein.^16^ A growing set of fluorescent sterol derivatives is now commercially available, which are often used interchangeably in the field. However, almost none of these probes have been profiled with regards to their selectivity towards sterol-binding proteins. This can lead to erroneous or incomplete interpretation of biological results generated with these probes. Rectifying this would also enable a more targeted and specific use of these probes to assess the biology of individual or groups of sterol-binding proteins. Recently we reported the initial stages of development of a sterol transport protein screening panel as a tool for the identification and characterization of potent and selective STP inhibitors. The screening platform, as such, enabled the identification of potent and selective OSBP binders, inhibiting retrograde trafficking and reducing Shiga toxin toxicity.^17^

Here we describe the profiling of fluorescent sterol-based probes as well as a set of steroid-based natural products, utilizing the now more comprehensive STP screening panel, providing the molecular basis for ligand recognition of ten STPs for the first time combined in one study. The identification of suitable fluorescent probes for the differential STPs enabled the establishment of fluorescence-, FRET-, and differential scanning fluorimetry-based assays, facilitating the direct comparison of STP specific selectivity profiles and evaluation of selective STP inhibitors in a high throughput manner. The screening of 41 sterol- and steroid-based natural products revealed Aster-A and Aster-B as targets for endogenous and synthetic steroid hormones and their precursors as well as STARD5 as the primary target for bile acid derivatives. Finally, those results combined with a detailed structural analysis revealed key residues, which could serve as selectivity handles for the development of highly potent and selective STP inhibitors in the future.

## RESULTS

### Sterol binding domains show differential binding affinity towards fluorescent sterol-based probes

We initially sought to assess the binding selectivity of different fluorescent sterols against STPs. To do this, we employed our recently developed STP screening panel comprising OSBP, ORP1 and ORP2 as members of the ORP family, STARD1, as well as Aster-A, Aster-B and Aster-C. To further expand this screening platform, we expressed and purified the sterol binding domains of STARD3, STARD4 and STARD5 harboring an N-terminal His_6_ tag. Circular dichroism spectroscopy confirmed the correct folding of all ten STPs (for CD spectra, please see Supplementary Figure 1). We screened six different sterol-based fluorophores against each sterol-binding domain (SBD, Figure 1a). Their fluorescence properties (excitation and emission spectra, Supplementary Figure 2) enabled the determination of dissociation constants (k_d_) by direct titration against increasing protein concentrations and monitoring changes in fluorescence intensity or fluorescence polarization (Table 1).

**Figure 1:**
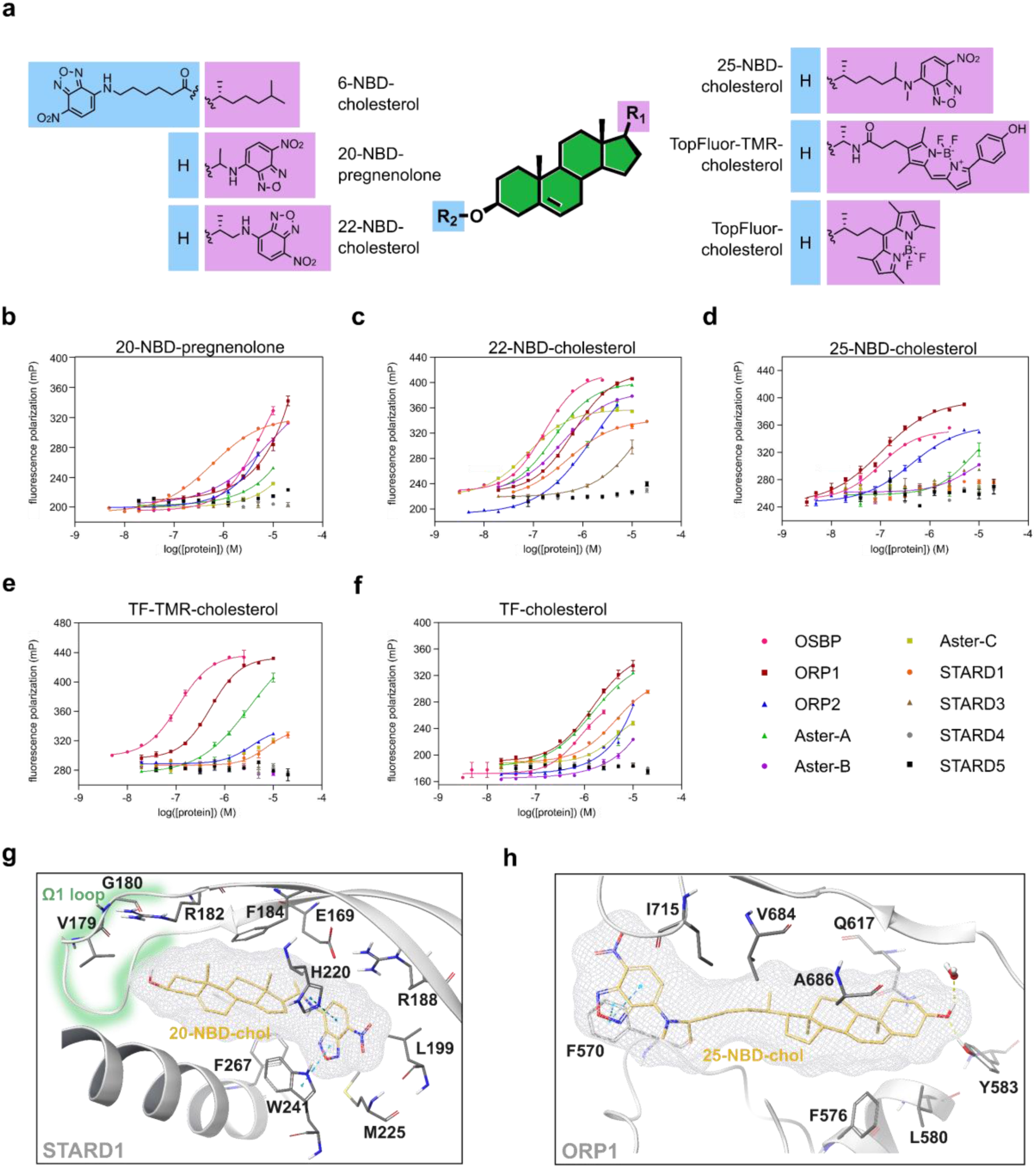
Investigation of sterol-based fluorescent probes towards their binding affinity against sterol transport proteins. **a** Representation of the chemical structures of the sterol-based NBD-labeled fluorescent probes. Direct titration of sterol-based fluorescent probes against increasing protein concentrations enables the determination of kd by monitoring changes in fluorescence polarization for **b** 20-NBD-pregnenolone, **c** 22-NBD-cholesterol, **d** 25-NBD-cholesterol, **e** TopFluor-TMR-cholesterol and **f** TopFluor-cholesterol. Experimental points were measured in duplicates on each plate and were replicated in n = 3 biologically independent experiments. Error bars indicate s.e.m. **g** Predicted binding pose of 20-NBD-pregnenolone into the crystal structure of STARD1 (pdb: 3p0l) indicating possible interactions with W24 and H220. **h** Predicted binding pose of 25-NBD-chol into the ORP1 crystal structure (pdb: 5zm5) highlighting possible interactions with Y583 and F570.

**Table 1:** Summary of binding affinities of sterol-based fluorescent probes against ten different sterol transport proteins. FP = fluorescence polarization; na = not applicable. All data is the mean of three biologically independent experiments.

|  | OSBP | ORP1 | ORP2 | Aster-A | Aster-B | Aster-C | STARD1 | STARD3 | STARD4 | STARD5 |
| --- | --- | --- | --- | --- | --- | --- | --- | --- | --- | --- |
| | FP - $K_d$<br>[nM] | FP - $K_d$<br>[nM] | FP - $K_d$<br>[nM] | FP - $K_d$<br>[nM] | FP - $K_d$<br>[nM] | FP - $K_d$<br>[nM] | FP - $K_d$<br>[nM] | FP - $K_d$<br>[nM] | FP - $K_d$<br>[nM] | FP - $K_d$<br>[nM] |
| 6-NBD-choI | > 5000 | > 5000 | 4560 | > 5000 | > 5000 | > 5000 | > 5000 | > 5000 | > 5000 | > 5000 |
| 20-NBD-preg | 4042 | > 5000 | > 5000 | > 5000 | 4818 | > 5000 | 487 | > 5000 | > 5000 | > 5000 |
| 22-NBD-choI | 149 | 322 | 4648 | 357 | 388 | 63 | 573 | > 5000 | > 5000 | > 5000 |
| 25-NBD-choI | 105 | 114 | 588 | > 5000 | > 5000 | > 5000 | > 5000 | > 5000 | > 5000 | > 5000 |
| TF-TMR-choI | 132 | 609 | 2920 | 3262 | > 5000 | > 5000 | > 5000 | > 5000 | > 5000 | > 5000 |
| TF-choI | 559 | 1162 | > 5000 | 1472 | > 5000 | 4805 | 4803 | > 5000 | > 5000 | > 5000 |

6-NBD-cholesterol (6-NBD-chol) shows weak or no binding of the STPs (Supplementary Figure 3), most likely due to the fact that sterols are predicted to bind “head-first” to all STPs, and thus substitution on the A-ring is not tolerated. 20-NBD-pregnenolone (20-NBD-preg) only binds STARD1 tightly, with a >10-fold selectivity over other STPs (Figure 1b). Furthermore, we observed a STARD1 specific increase in fluorescence upon 20-NBD-preg binding (Supplementary Figure 4), which was not the case for other NBD-chol probes, suggesting a distinct binding mode (*vide infra*).^18^ 22-NBD-cholesterol (22-NBD-chol) appears to be the most universal probe, showing a k_d_ below 500 nM for six out of ten STPs (Figure 1c). Most noticeably, it does not show any binding affinity towards STARD3/4/5. On the other hand, 25-NBD-cholesterol (25-NBD-chol), which harbors a longer linker between the cholesterol core and NBD-group, shows selective binding to the ORP family proteins OSBP, ORP1 and ORP2 (Figure 1d). TF-TMR-cholesterol (TF-TMR-chol) shows preferred binding to OSBP (k_d_ = 132 nM) while having a lower binding affinity towards the other STPs, suggesting a potential utility as a selective fluorescent probe (Figure 1e). This result is consistent with the recently reported importance of an amide linker between the sterol core and sidechain, which is suggested to interact with Thr491, conferring tight OSBP binding and high selectivity.^17^ Furthermore, TF-TMR-chol could serve as an alternative probe to the NBD-labeled probes for different purposes, since it is excited and emits at different wavelengths (Supplementary Figure 2).

In contrast, TF-chol shows an overall weaker binding to the STPs suggesting that a longer linker between the cholesterol core and the BODIPY group might be preferred. However, STARD3, STARD4 and STARD5 don’t bind any of the side chain labeled fluorescent probes, suggesting that they might prefer much smaller ligands because of their short and narrow binding pockets. While crystal structures of several STPs have been reported to date, only OSBP, ORP1 as well as murine Aster-A and Aster-C have been crystallized in complex with a ligand binding in the sterol binding pocket. To obtain insights into the specific binding modes of selected fluorophores and to rationalize their biological effect we performed docking studies utilizing an induced fit-based approach. Interestingly, the most conserved poses of 20-NBD-preg docked into the crystal structure of STARD1 (pdb: 3p0l^19^) with a head-out orientation, where the NBD-group binds deep into the sterol binding pocket. The modeling predicts interactions of the NBD-group with H220 and W27 providing a plausible explanation for its turn-on fluorescence upon binding to STARD1 (Figure 1g). The modeling of 25-NBD-chol into the ORP1 crystal structure (pdb: 5zm5^20^) suggest a head-in orientation with the hydroxy-group interacting with a water molecule as well as Y583. Additionally, the NBD-group is predicted to interact with F570 located at the opening of the sterol binding pocket, which is conserved among the ORPs (OSBP F440; ORP2 F69) and thereby explaining their tight binding to 25-NBD-chol (Figure 1h).

### Biophysical thermal shift assay and sterol transfer assay reveal 25-HCTL and 27-HCTL as potent STARD protein inhibitors

Due to the lack of a suitable fluorescent probe for the development of a competitive fluorescence-based assay for STARD3, STARD4 and STARD5, we investigated differential scanning fluorimetry (DSF). In previous studies this method has already proven to be useful for the screening of potent and selective Aster inhibitors.^21,22^ Upon incubation of the STARD proteins with SYPRO Orange, we could observe usable melting curves for STARD3, STARD4 and STARD5 (Figure 2a). However, for STARD1 no melting curve was observed. Direct titration of SYPRO Orange against increasing concentrations of the individual STARD proteins revealed its binding affinity to STARD1 in the nanomolar range and thereby explaining interference with the DSF assay (Figure 2b). The same effect was observed for ORP1 and ORP2, however, not for Aster-A-C, thus proving the correlation between SYPRO Orange binding and the usability of DSF for the specific STP (Supplementary Figure 4). Next, we sought out to investigate intrinsically fluorescent sterol probes. As dehydroergosterol (DHE) had very poor solubility and would not be suitable as a tracer in fluorescence-based assays where it has not previously been integrated into membranes, we tested single concentrations of the intrinsically fluorescent oxysterols 25-hydroxycholestatrienol (25-HCTL) and 27-hydroxycholestatrienol (27-HCTL) (Figure 2c) against STARD3-5, as we predicted that their smaller size may be better tolerated by this class of STPs.^23,24^ We observed a strong stabilization of all three proteins by 27-HCTL, while 25-HCTL preferably stabilizes STARD3 and STARD4 (Figure 2d-f and Table 1). As an orthogonal assay, we employed variable temperature measurements using circular dichroism as a readout to confirm binding of 25-HCTL and 27-HCTL to the specific STPs (Supplementary information Figure 5). We expanded the profiling of 25- and 27-HCTL by screening them as competitive inhibitors in FP assays against the remaining STPs (Table 2). Interestingly, 27-HCTL seems to be more promiscuous towards STPs (Table 2), which could be explained by its longer and more flexible side chain, a result of the position of the hydroxyl group on the terminal methyl in the sterol side chain. This gave early indications that small structural changes can lead to differential selectivity profiles towards the STPs.

**Figure 2:**
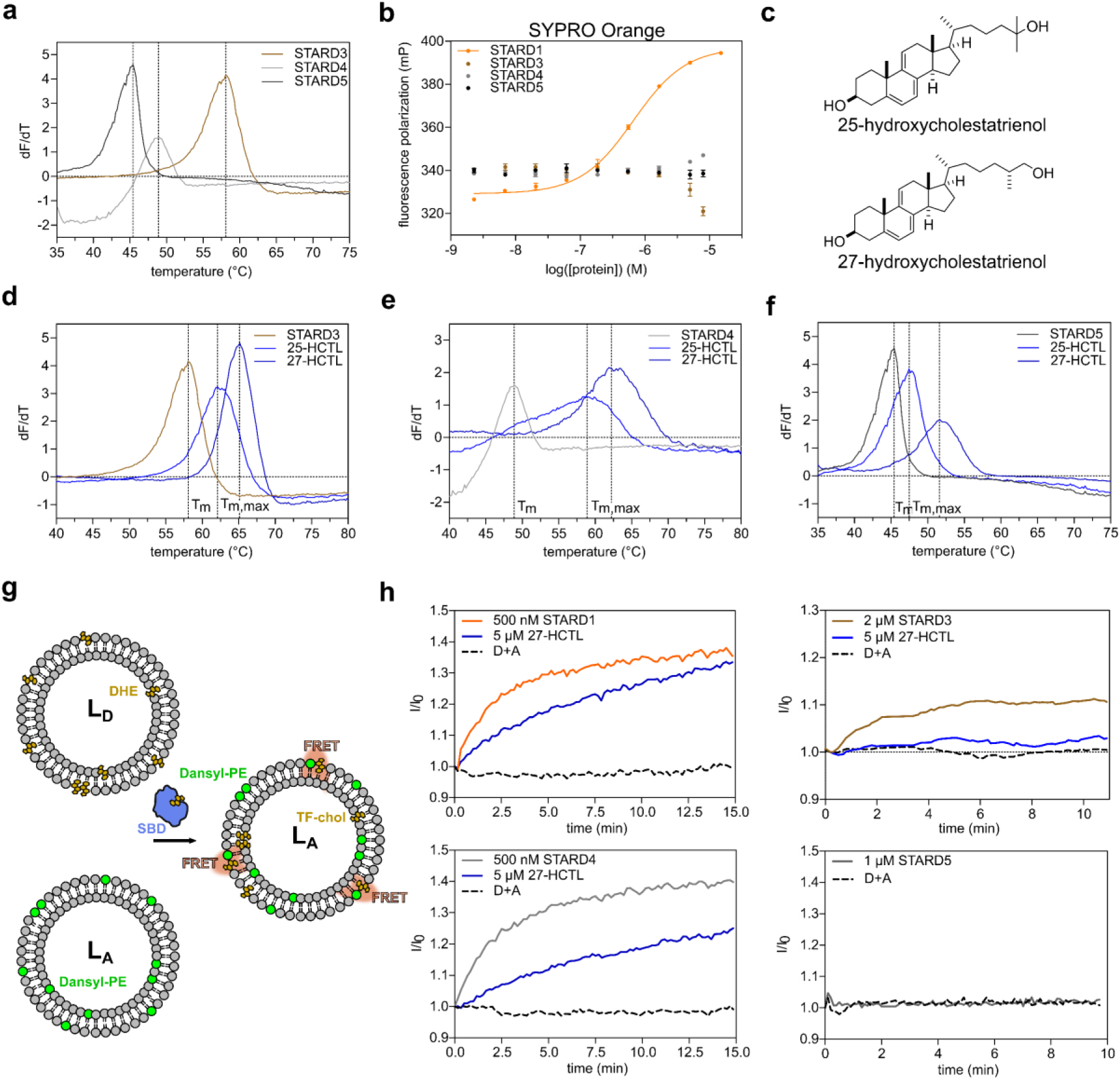
25-HCTL and 27-HCTL stabilize STARD3, STARD4 and STARD5. **a** Thermal stability of STARD3, STARD4 and STARD5 assessed by differential scanning fluorimetry. **b** Differences in fluorescence polarization upon titration of SYPRO Orange against increasing protein concentrations showing binding of STARD1 to SYPRO Orange. **c** Chemical structures of 25-hydroxycholestatrienol (25-HCTL) and 27-hydroxycholestatrienol (27-HCTL). **d** Thermal stabilization of STARD3 incubated with single concentrations of 25-HCTL and 27-HCTL. **e** Thermal stabilization of STARD4 incubated with single concentrations of 25-HCTL and 27-HCTL. **f** Thermal stabilization of STARD5 incubated with single concentrations of 25-HCTL and 27-HCTL. **g** Schematic of the FRET-based sterol transport assay used for the STARDs. **h** Transport of dehydroergosterol (DHE) by STARD1/3/4/5, and its inhibition by 27-HCTL, as assed by a FRET assay. D = Donor liposome, A = acceptor liposome. All data represents a representative experiment from three biological replicates (n = 3).

**Table 2:** Summary of the binding affinities of 25- and 27-HCTL measured by using a competitive FP assay as well as a thermal shift assay. FP = fluorescence polarization. ΔTm,max refers to the maximal stabilization of a protein by the compound across all concentrations. All data is the mean of three biologically independent experiments.

|  | OSBP | ORP1 | ORP2 | Aster-A | Aster-B | Aster-C | STARD1 | STARD3 | STARD4 | STARD5 |
| --- | --- | --- | --- | --- | --- | --- | --- | --- | --- | --- |
| | FP - IC<sub>50</sub><br>[nM] | FP - IC<sub>50</sub><br>[nM] | FP - IC<sub>50</sub><br>[nM] | FP - IC<sub>50</sub><br>[nM] | FP - IC<sub>50</sub><br>[nM] | FP - IC<sub>50</sub><br>[nM] | FP - IC<sub>50</sub><br>[nM] | $\Delta T_{m,max}$<br>[°C] | $\Delta T_{m,max}$<br>[°C] | $\Delta T_{m,max}$<br>[°C] |
| 25-HCTL | 238 | 1303 | 3115 | > 10000 | 6960 | 8255 | 3121 | 5.9 | 10.3 | 3.2 |
| 27-HCTL | 292 | 777 | 1983 | 625 | 2242 | 3060 | 2019 | 7.4 | 13.8 | 8.6 |

Next, we employed an in vitro assay based on Förster (fluorescence) resonance energy transfer (FRET) to evaluate the effect of 27-HCTL on the sterol transfer functions of the STARD proteins. Here, the transfer of the fluorescent cholesterol analogue DHE between a donor (L_D_) and an acceptor liposome (L_A_) is followed. The L_D_ liposomes contain DHE while the L_A_ contain Dansyl-PE. The STP-mediated transfer of DHE from the L_D_ to the L_A_ results in DHE and Dansyl-PE forming a FRET pair and thereby leading to an increase in fluorescence intensity signal (Figure 2g).^25^ Notably, we could observe a transport of DHE by STARD1, STARD3 and STARD4 only, while for STARD5 no increase in fluorescence was observed. Those results indicate that STARD5 might not be involved in cholesterol transport but in the transport of other ligands. We were able to observe a decreased FRET signal for STARD1, STARD3, and STARD4 incubated with 27-HCTL, indicating its inhibition of sterol transport.

### Intrinsic fluorescence of 25-HCTL and 27-HCTL reveals their high affinity binding to the STARDs

As no suitable fluorescent probe for STARD3, STARD4, and STARD5 was identified, the intrinsic fluorescence of 25-HCTL and 27-HCTL was investigated. UV-vis and fluorescence spectra revealed excitation peaks around 325 nm and emission peaks around 420 nm. The comparison between spectral data of 25- and 27-HCTL measured in buffer, methanol and DCM showed sensitivity to the environment resulting in changes in intensity and shifts in emission wavelength (Figure 3a-b). Following this, we hypothesized that binding into a hydrophobic sterol binding domain could result in changes in the emission spectra. Upon titration of increasing SBD concentrations, we observed an increase in fluorescence for all four STARD proteins (Supplementary Figure 7a-b). To confirm the specificity of this effect we employed FRET measurements between the protein’s tryptophan residues and the intrinsically fluorescent sterol, as previously described to confirm specific binding of macarangin B enantiomers to OSBP.^26^ Using an excitation wavelength of 280 nm and an emission wavelength of 450 nm we were able to measure the specific k_d_’s of 25-HCTL and 27-HCTL for the STARD proteins (Table 3 and Figure 3c-d). The results correlate with the previously measured IC_50_ (STARD1) and the ΔT_m,max_ values (STARD3, STARD4, STARD5) showing tight binding of all four STARD proteins to 27-HCTL, while 25-HCTL binds STARD1, STARD3 and STARD4 only (Figure 3e). Based on these results we investigated the usability of both ligands as fluorescent probes for the development of competitive binding assays by monitoring changes in FRET-based fluorescence intensity. While 80 nM probe incubated with 120 nM STARD4 gave a sufficient assay window and a Z-factor of 0.8, we couldn’t observe sufficient windows for the other STARD proteins (Figure 3g and Supplementary Figure 7c-d) suggesting that 25- and 27-HCTL are suitable fluorescent probes for STARD4 only. The use of higher protein concentrations did not improve the assay window. Modeling of 25-HCTL into the STARD4 crystal structure (pdb: 6l1d)^25^ revealed a possible explanation for the much higher increase in fluorescence intensity upon binding to STARD4 in comparison to the other STARD proteins (Figure 3f). The most conserved poses predict a head-in conformation of 25-HCTL in the sterol binding pocket and close-proximity of the ligand to W155, promoting direct FRET between ligand and protein.

**Figure 3:**
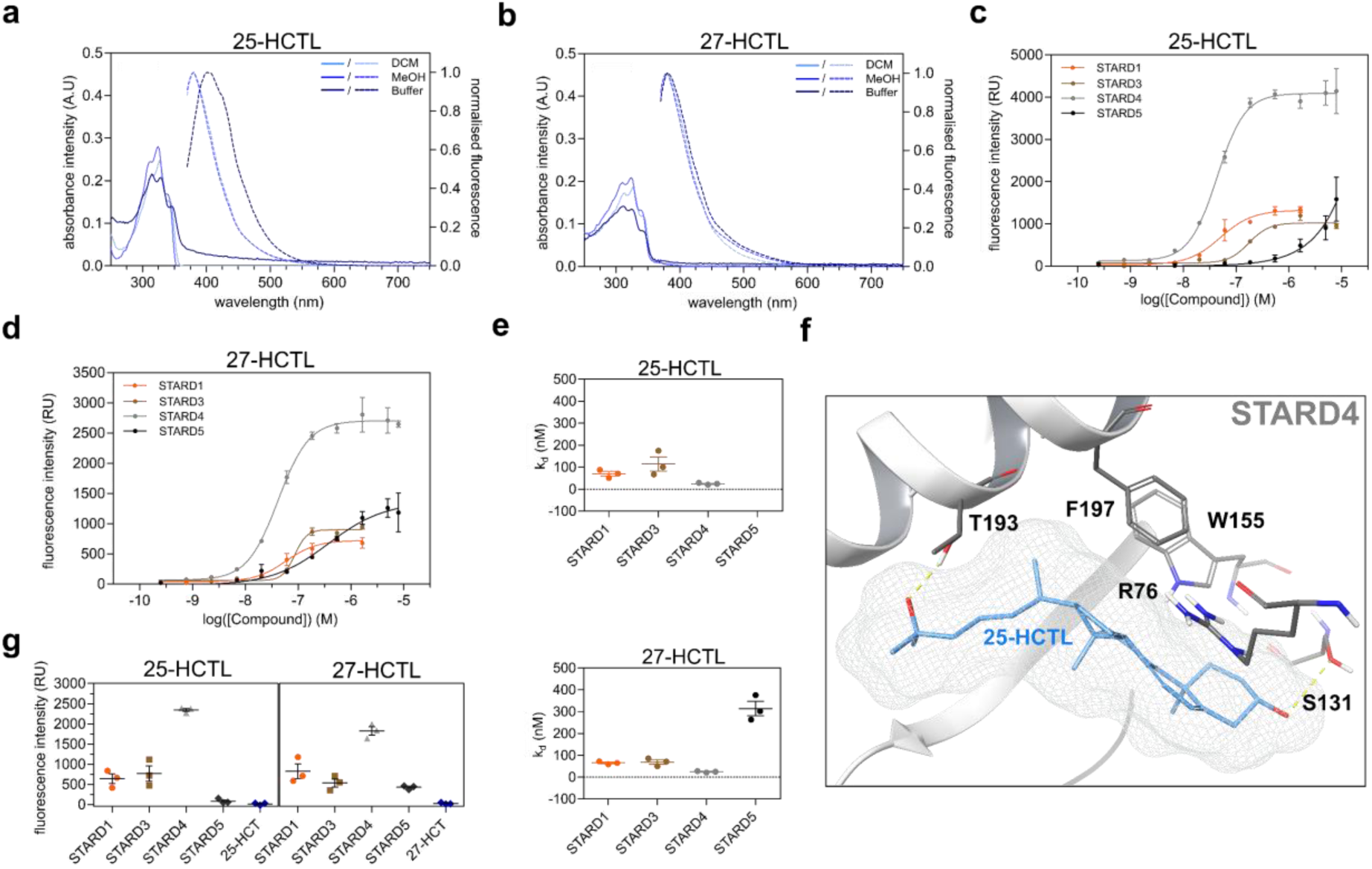
Inherent fluorescence of 25- and 27-HCTL enables the determination of dissociation constants. **a** Excitation and emission (excitation at 325 nM) spectra of 25-HCTL in DCM, methanol or HEPES buffer. **b** Excitation and emission (excitation at 325 nM) spectra of 27-HCTL in DCM, methanol or HEPES buffer. **c** Titration of 25-HCTL against STARD1, STARD3, STARD4 and STARD5 assessed by FI using an excitation wavelength of 280 nm and an emission wavelength of 450 nm. **d** Titration of 27-HCTL against STARD1, STARD3, STARD4 and STARD5 assessed by FI using an excitation wavelength of 280 nm and an emission wavelength of 450 nm. **e** Dissociation constants of 25- and 27-HCTL for STARD1, STARD3, STARD4 and STARD5**. f** Representation of the most conserved binding pose of 25-HCTL docked into the crystal structure of STARD4 (pdb: 6l1d**). g** Representation of fluorescence intensity windows for 80 nM 25-HCTL and 27-HCTL incubated with the different STARD proteins.

**Table 3:** Summary of the binding affinities of selected STP hits measured by using a competitive FP/FI assay as well as a thermal shift assay. FP = fluorescence polarization. DSF = differential scanning fluorimetry. All data is the mean of three biologically independent experiments. All ligands show an IC50 > 10000 nM for STARD4 as assessed by FI and were therefore removed from the table. The definition of all compound abbreviations can be found in Supplementary Table 1.

|  | OSBP | ORP1 | ORP2 | Aster-A | Aster-B | Aster-C | STARD1 | STARD3 | STARD5 |
| --- | --- | --- | --- | --- | --- | --- | --- | --- | --- |
|  | FP - IC<sub>50</sub><br>[nM] | FP - IC<sub>50</sub><br>[nM] | FP - IC<sub>50</sub><br>[nM] | FP - IC<sub>50</sub><br>[nM] | FP - IC<sub>50</sub><br>[nM] | FP - IC<sub>50</sub><br>[nM] | FP - IC<sub>50</sub><br>[nM] | DSF -<br>ΔT<sub>m,max</sub><br>[°C] | DSF -<br>ΔT<sub>m,max</sub><br>[°C] |
|  | competitive<br>22-NBD-chol | competitive<br>25-NBD-chol | competitive<br>25-NBD-chol | competitive<br>22-NBD-chol | competitive<br>22-NBD-chol | competitive<br>22-NBD-chol | competitive<br>22-NBD-chol |  |  |
| 4β-HC | 1684 | > 10000 | > 10000 | > 10000 | > 10000 | > 10000 | > 10000 |  |  |
| 7α-HC | 361 | 9234 | > 10000 | > 10000 | > 10000 | > 10000 | > 10000 |  |  |
| 7β-HC | 838 | > 10000 | > 10000 | > 10000 | > 10000 | > 10000 | > 10000 |  |  |
| 7-KC | 189 | 1578 | 1154 | 306 | 2482 | 4418 | > 10000 |  |  |
| 20(*S*)-HC | 41 | 1190 | 1308 | 1731 | 721 | 2839 | 4472 |  |  |
| 22(*R*)-HC | > 10000 | 8022 | 2853 | 9187 | 1582 | > 10000 | > 10000 |  |  |
| 24(*S*)-HC | 213 | 3332 | 2541 | > 10000 | 7619 | > 10000 | 6205 |  |  |
| 25-HC | 13 | 882 | 1159 | 622 | 1895 | > 10000 | 997 | 2.2 | 1.3 |
| 27-HC | 124 | 1062 | 2736 | > 10000 | 3369 | > 10000 | 4207 | 3.5 | 1.2 |
| CT | 6192 | > 10000 | > 10000 | > 10000 | > 10000 | > 10000 | > 10000 |  |  |
| CA | > 10000 | > 10000 | > 10000 | > 10000 | > 10000 | > 10000 | > 10000 |  | 5.0 |
| CDCA | > 10000 | > 10000 | > 10000 | > 10000 | > 10000 | > 10000 | > 10000 | 0.5 | 3.3 |
| DCA | > 10000 | > 10000 | > 10000 | > 10000 | > 10000 | > 10000 | > 10000 |  | 7.5 |
| HDCA | > 10000 | > 10000 | > 10000 | > 10000 | > 10000 | > 10000 | > 10000 |  | 3.7 |
| AD | > 10000 | > 10000 | > 10000 | > 10000 | 2704 | > 10000 | > 10000 |  |  |
| DHEA | > 10000 | > 10000 | > 10000 | 672 | 656 | > 10000 | > 10000 |  |  |
| PRG | > 10000 | > 10000 | > 10000 | 417 | 822 | > 10000 | > 10000 |  |  |
| 21-AcP | > 10000 | > 10000 | > 10000 | 1932 | 801 | > 10000 | > 10000 | 2.5 | 1.3 |
| 3-AcP | 999 | 7258 | > 10000 | > 10000 | 3941 | > 10000 | 6060 |  |  |
| DHEA-Ac | 5529 | > 10000 | 9315 | > 10000 | > 10000 | > 10000 | > 10000 |  |  |
| U18666A | > 10000 | > 10000 | 7817 | 2319 | 931 | 3537 | > 10000 |  |  |
| T | > 10000 | > 10000 | > 10000 | > 10000 | > 10000 | > 10000 | > 10000 | 1.6 | 1.1 |
| Me-DHT | > 10000 | > 10000 | > 10000 | > 10000 | > 10000 | > 10000 | > 10000 |  |  |
| PG | > 10000 | > 10000 | > 10000 | > 10000 | 1971 | > 10000 | > 10000 | 3.3 | 3.5 |
| TP | > 10000 | 4415 | 7160 | > 10000 | 762 | > 10000 | 6527 | 4.3 | 3.0 |
| 6-DHT-Ac | > 10000 | 9996 | > 10000 | > 10000 | 1486 | > 10000 | > 10000 | 4.5 | 3.7 |

### Biophysical screening platform reveals differential selectivity profiles of sterol transport proteins

To gain a deeper insight into the individual selectivity profiles of the STPs we employed the biophysical screening platform and screened a set of 41 natural products, which can be further divided into 4 different compound classes. We tested 9 oxysterols, 26 steroid hormones and hormone precursors, 5 cholic acid derivatives as well as the phytosterol β-sitosterol. While NBD-labeled probes were used for the ORPs, the Asters and STARD1, we used 27-HCTL as a fluorescent probe for STARD4. For STARD3, STARD4, and STARD5 we used DSF as an alternative and orthogonal assay. First, a single concentration (10 µM) screen was performed to identify potential STP binders (Supplementary Dataset 1). We chose 50% inhibition for the competitive assays and a ΔT_m_ of two degrees for the DSF assay as a cut-off for further investigations. Compounds that fulfilled these criteria were tested in dose response against all ten STPs (Table 3). Interestingly, in this compound library we couldn’t identify STARD4 ligands with affinities <10 µM. However, the results show that a differential hydroxylation pattern on the cholesterol core can result in differential selectivity profiles towards the STPs. OSBP binds most of the oxysterols with high affinity. 4*β*-HC, 7*ɑ*-HC and 7*β*-HC selectively bind to OSBP showing that hydroxylation’s at the sterol core are more tolerated in OSBP than the other STPs.^27^ Furthermore, 20(*S*)-HC binds OSBP with a k_d_ of 41 nM showing a > 20-fold selectivity ratio towards the other STPs. Retrospectively, the high affinity and selectivity of 20(*S*)-HC to OSBP may provide an explanation for the Golgi accumulation of a fluorescent alkynyl derivative of 20(*S*)-HC observed in previous studies.^28^ In contrast, side chain-modified oxysterols show promiscuous binding to several STPs. However, OSBP does not show binding to 22(*R*)-HC while ORP1, ORP2 as well as Aster-B do, suggesting an unfavorable positioning of the OH-group in the OSBP binding pocket.^29^ An overlay of the predicted poses of 22(*R*)-HC and 25-HC in the recently published crystal structure of OSBP (pdb: 7v62)^30^ reveals an explanation for their differential binding affinities (Supplementary Figure 8b). While both ligands are predicted to bind in the “head-in” conformation, 25-HC possibly interacts with T491 as well as D453 and K577. The hydroxylation at position 22 in the (*R*)-configuration seems to result in a different orientation of the sterol core hindering favorable interactions with the key residues in the OSBP binding pocket.

In previous studies, contradictory results regarding the binding preference of STARD5 were suggested. While some groups report the binding of cholesterol and some oxysterols, other groups described STARD5 as a bile acid binder.^31,32^ To obtain further insights into STARD5’s binding preferences, we employed our STP screening panel. The results suggest exclusive binding of cholic acid derivatives to STARD5, with DCA displaying the highest ΔT_m,max_ (7.5 °C) (Table 3). Furthermore, no other STP in our assay panel bound cholic acid derivatives, suggesting a specific function of STARD5 in bile acid distribution. Docking studies revealed a distinct interaction pattern between STARD5 (pdb: 2r55)^19^ and cholic acid. While the 3-OH-group is predicted to interact with V68 and the carboxylic acid with T103, the OH-groups on the B-and C-ring interact with R76. The nonplanar shape of CA due to its *cis* A/B ring fusion seems to orient the molecule in a favorable distance to R76 and the backbone of V68. This might result in an unfavorable orientation for ligands with more planar conformations, providing a plausible explanation for weak (25-HC and 27-HC) and no binding of other Δ^5^-oxysterols to STARD5 (Supplementary Figure 9).

Steroidogenesis is a highly complex multienzyme process converting cholesterol into biologically active steroid hormones, which is largely confined to the adrenal cortex, testicular Leydig cells, ovarian granulosa and theca cells.^33^ Some STPs including Aster-B as well as STARD1/3/4/5 are highly expressed in steroidogenic tissues.^34^ To gain further insight into the binding capabilities of STPs to endogenous and synthetic steroid hormones and their precursors we evaluated a set of 26 ligands (Table 3). STARD1 transfers cholesterol from the OMM to the IMM, initiating its conversion to pregnenolone, and thereby mediating an acute steroidogenic response.^35^ However, none of the tested ligands bind STARD1 with an IC_50_ <5 µM suggesting that it is not involved in the direct transfer of those steroid hormones and their precursors. It is thus also unlikely that STARD1 participates in a negative feedback loop where steroidogenic products inhibit their own synthesis by limiting cholesterol availability. In previous studies, 21-Acetoxypregnenolone (21-AcP) was shown to inhibit steroid synthesis in murine MA-10 Leydig tumor cells, with STARD1 as the proposed target.^36^ Surprisingly, our data suggests that Aster B, and possibly Aster-A, are the primary targets for 21-AcP. Our STP screening panel revealed Aster-A and Aster-B’s equipotent binding of dehydroepiandrosterone (DHEA) (Figure 4b), pregnenolone (PRG) and 21-AcP all harboring a 3-OH-group as well as a 5,6-alkene in the B-ring (Figure 4a). Acetylation of the 3-OH-group thereby seems to be less tolerated although alkylation is tolerated as exemplified by the pan-Aster inhibitor U18666A.^37^ Interestingly, there is stereospecificity in the binding to the Asters as the enantiomer of U18666A was inactive against all STPs in the panel.^38^ Progesterone (PG), testosterone propionate (TP) (Figure 4c) and 6-dehydrotestosterone acetate (6-DHT-Ac) strongly bind Aster-B only, harboring a 3,4,5-*ɑ,β*-unsaturated ketone in the A-ring (Figure 4a).

**Figure 4:**
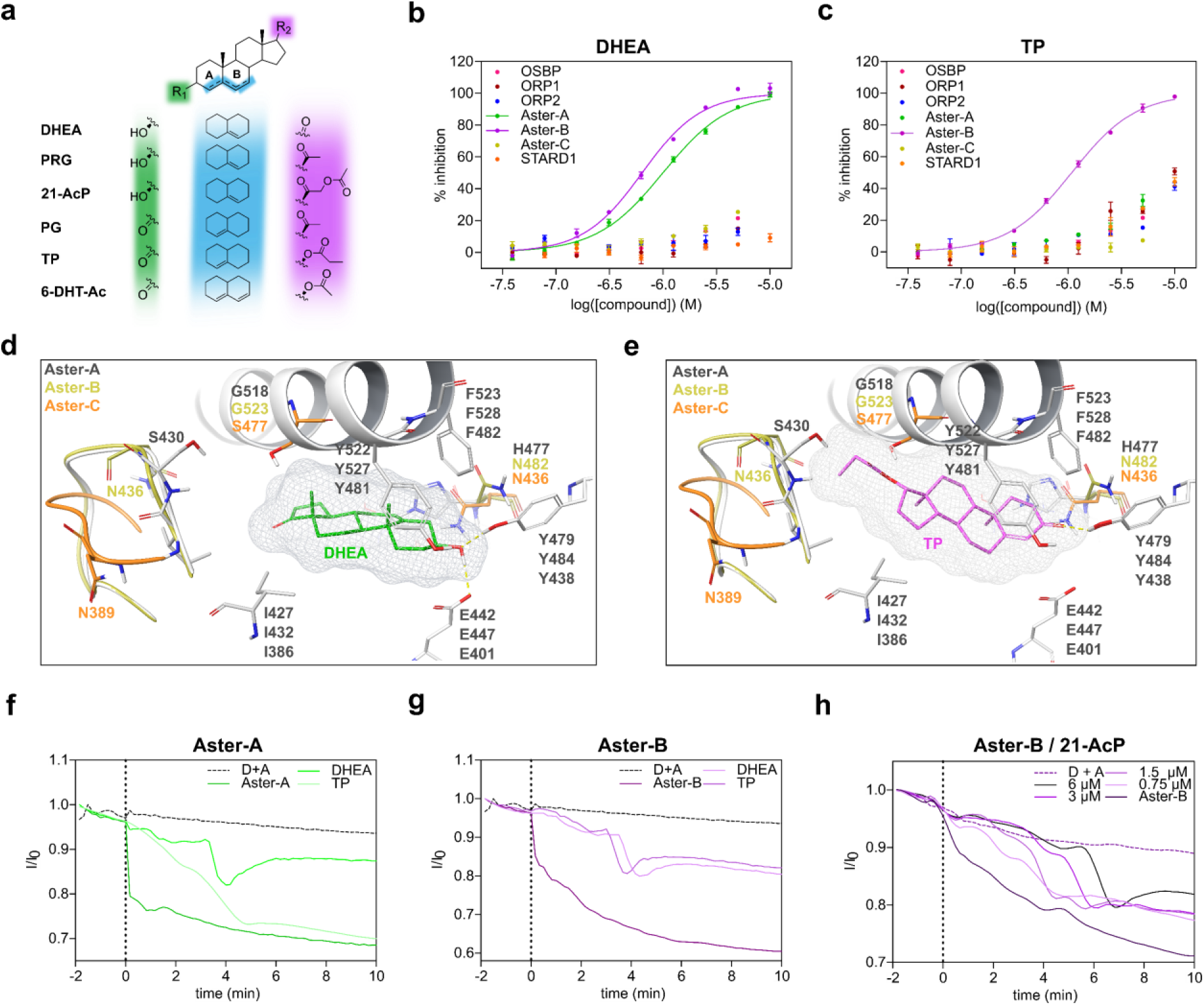
Biophysical screening platform reveals binding of steroid hormone precursors to Aster-B. **a** Chemical structure of the Aster-A and Aster-B binding endogenous and synthetic steroid hormones. **b** Dose-dependent inhibition of STPs binding to NBD-chol by DHEA as assessed by FP. One representative experiment is shown from three independent experiments (n=3). **c** Dose-dependent inhibition of STPs binding to NBD-chol by TP as assessed by FP. One representative experiment is shown from three independent experiments (n=3). **d** Representation of the most conserved binding pose of DHEA docked into the crystal structures and homology models of the Asters. **e** Representation of the most conserved binding pose of TP docked into the crystal structures and homology models of the Asters. **f** Inhibition of sterol transport mediated by Aster-A (125 nM, left) by DHEA and TP. One representative experiment is shown from two independent experiments (n=2), D = donor liposomes, A = acceptor liposomes. **g** Inhibition of sterol transport mediated by Aster-B (125 nM, left) by DHEA and TP. One representative experiment is shown from two independent experiments (n=2), D = donor liposomes, A = acceptor liposomes. **h** Inhibition of sterol transport mediated by Aster-B (125 nM, left) by different concentrations of 21-AcP. One representative experiment is shown from two independent experiments (n=2), D = donor liposomes, A = acceptor liposomes.

When comparing the sterol binding pockets of Aster-A, -B and -C, only a few differences can be observed. To rationalize this selectivity profile, we modeled DHEA as well as TP into the Aster binding pockets (Aster-A pdb: 6gqf; Aster-B homology model based on 6gqf; Aster-C pdb: 7azn; Figure 4d-e).^13,37^ Whereas DHEA forms key interactions with E442/447 and Y479/484 and perfectly embeds into the Aster-A and B binding pocket (Figure 4c), TP forms interactions with Y484 only and the sterol core is significantly rotated, inducing a possible steric clash with H477 in Aster-A. Furthermore, Aster-C contains a serine (S477) in the middle of the binding pocket instead of a more spacious glycine in Aster-A (G518) and Aster-B (G523), suggesting a steric clash with the sterol core as a plausible explanation for its weak to no binding of the tested ligands. Next, we investigated the effect of DHEA and TP on the Aster-A and Aster-B mediated transport of 23-BODIPY-cholesterol (TF-chol) between artificial liposomes, by monitoring changes in FRET fluorescence intensity signal with rhodamine 1,2-dihexadecanoyl-sn-glycero-3-phospho-ethanolamine (Rh-DHPE) (Figure 4f-h). While Aster- A-mediated TF-chol transport is only inhibited by DHEA (Figure 4f), Aster-B-mediated transport is inhibited by DHEA and TP (Figure 4g) confirming the differential binding preferences of those ligands. Interestingly, despite using a large excess of ligand no full inhibition of transport could be achieved. Furthermore, we also observed the dose-dependent inhibition of Aster-B-mediated TF-chol transfer by 21-AcP (Figure 4h). Interestingly, for all investigations into the inhibition of Aster-mediated sterol transfer, there is a dose dependent lag-time from the point at which protein-ligand addition occurs and when the rate of transfer increases steeply. As the competing steroid is present in excess, the binding to the STP is initially saturated, and no transport of TF-chol is observed. However, due to the possible integration of the steroid into the membrane, the transfer of TF-chol proceeds quickly once the competing steroid has been integrated. As such, the data may reflect two processes: a) ligand-protein inhibition in solvent and b) ligand extraction from the membrane. This is further supported by reports showing that sterols and steroidal compounds including PRG, PG and DHEA can be embedded into lipid bilayers.^39–41^ To test this theory, two systems were compared: 1) as outlined above, the protein and ligand were pre-incubated for two minutes and then added to the liposome mixture; and 2) the ligand was preincubated with the liposomes for two minutes, followed by the addition of protein. In the second approach, unlike the first, the sterol transfer is initiated nearly immediately, however, similarly to the first, the maximal inhibition never approaches 100%. This test supports the idea that ligands both act as inhibitors of endogenous sterol transfer, as well as substrates directly from the liposome.

## DISCUSSION

Small molecules and natural products are widely used as powerful tools to explore the specific functions of proteins in the human cell. However, using poorly characterized and non-selective probes can lead to misleading and inconclusive results. In the last decades, a growing set of fluorescent and non-fluorescent sterol derivatives for studying sterol transport has become available. Those probes were extensively studied regarding their localization, interaction and trafficking in cells.^42–44^ However, the majority of those tool compounds have not been profiled with regards to their selectivity towards sterol-binding proteins, leading to inaccurate or incomplete interpretation of biological results. Therefore, we set out to exploit our STP screening panel to characterize different fluorescent sterols regarding their binding and selectivity profiles towards ten STPs. We found that especially the labeling-position as well as the linker properties are critical for the observed binding preferences. For example, the widely used 22-NBD-chol binds the majority of STPs in our screening panel, whereas 25-NBD-chol selectively binds to the ORP family proteins. In contrast to that, TF-TMR-chol harboring an amide in the linker has a 6-fold lower k_d_ towards OSBP in comparison to ORP1 and a 30-fold lower k_d_ in comparison to ORP2 and is inactive on the other STPs. We show that it is highly important to carefully choose the probe and the applied concentration for a specific research question. For this, we envisage that the data provided herein will be a valuable resource to guide and facilitate the interpretation of biological results within the field. Furthermore, we identified the inherent fluorescent sterols 25-HCTL and 27-HCTL as potent inhibitors of STARD3, STARD4 and STARD5 and were able to exploit 27-HCTL for the development of a fluorescence intensity-based competitive assay, which will facilitate the high-throughput screening for potent and selective STARD4 inhibitors in the future. However, with insufficient assay windows and Z’-factors 25- and 27-HCTL are not suitable for competitive fluorescence-based assays for STARD3 and STARD5, suggesting that the observed increase in fluorescence upon binding is highly dependent on the specific interaction between compound and protein. Here, DSF serves as an alternative method to detect and evaluate binding to STARD3 and STARD5.

Within the last decades, various studies reported binding data of different steroids to STPs, but information is provided across a diverse range of sources, thus making it difficult and tedious to find, evaluate and compare results. By developing a comprehensive set of biophysical tools, we were able to test a set of 41 steroid-based natural products and derivatives. Those results are complemented by a thorough docking-based structural analysis, identifying crucial amino acids in the STP binding pockets as molecular basis for their individual ligand selectivity. In general, it should be noted that due to the properties of the specific STPs and their differential binding affinities to the fluorescent probes, different protein concentrations had to be used in the different assay systems, influencing the individual assay limits and sensitivities. Nevertheless, our results prove that small changes on the steroid core, like a differential hydroxylation pattern, can result in varying binding affinities towards the STPs even within the same protein family. For example, OSBP tolerates hydroxylation directly at the steroid core as well as side-chain hydroxylation, while the other STPs mainly tolerate side-chain hydroxylation. Furthermore, our results confirm STARD5 as a bile acid binding protein, which could be explained by the presence of an Arg in the center of the pocket. Additionally, we were able to provide a plausible link between the inhibition of steroid synthesis in murine MA-10 Leydig tumor cells by identifying Aster-A and Aster-B as primary targets for 21-AcP, which was previously suggested to target STARD1. In particular, Aster-B is highly expressed in steroidogenic tissues. By identifying binding of DHEA, PRG and PG as well as their synthetic analogues 21-AcP, 6-DHT-Ac and TP to Aster-B our results suggest that Aster-B might be directly involved in steroidogenesis. This is supported by recent reports which suggest that Aster-B mediated cholesterol transport from the PM to the ER and from the ER to mitochondria directly regulates estradiol production.^12,45^

In summary, we established a comprehensive set of biophysical assays complemented by robust docking workflows for the evaluation of ligand selectivity towards sterol transport proteins. We utilized those tools to obtain unique insights into the selectivity profiles of 41 steroid-based natural products as well as the underlying molecular basis. In the future, this study will serve as a resource to guide the selection of suitable fluorescent probes for specific research questions and will guide and facilitate the interpretation of cell biological results. Furthermore, the insights into the STP binding pockets will guide the development of potent and selective probes for the investigation of STP specific functions in health and disease.

## Acknowledgements

The Laraia Laboratory was supported by funding from the Novo Nordisk Foundation (grant nos. NNF19OC0055818, NNF19OC0058183, NNF21OC0067188), the Carlsberg Foundation (grant no. CF19-0072) and the European Union (ERC, ChemBioChol, 101041783). We thank Mjaftime Ismaili and David Frej Nielsen for their support with the expression and purification of proteins. D.F.C was funded by Grant 1 P50 MH122379 from the National Institutes of Health.

## Author contributions

L.D. and L.L. designed the project. L.D. expressed and purified recombinant proteins and developed all biophysical assays. H.P.B-R. and N.C. performed fluorescence absorption and emission scans. L.D. and H.P.B-R. established and performed fluorescence intensity and FRET assays. L.D. A.W.B. and H.P.B-R. developed differential scanning fluorimetry assays. H.P.B-R., L.D. and N.J.D. performed and analyzed docking experiments and performed FP experiments. N.J.D. synthesized 20-NBD-pregnenolone. M.Q. and Y.X. synthesized 25- and 27-HCTL under supervision of D.F.C. L.L. supervised the project. L.D. and L.L wrote the paper with input from all authors.

## Competing interests

The authors declare no competing interest.

## MATERIALS AND METHODS

### Chemicals

Lipids were commercially available or synthesized in house (see supporting information for synthetic procedures). Information about supplier, catalogue number and CAS number are available in Supplementary table 1. Nuclear magnetic resonance spectra were analyzed with MestreNova, v.x64.

### Protein expression constructs

Human ASTER domains of Aster-A(359-547), -B(364-552) and -C(318-504) were subcloned into a pGEX-6p-2rbs vector, thus introducing the cloning artifact ‘GPLGS’.^15^ The pGEX-6P-1-GST-OSBP(377-807), pET24b(+)-ORP1(534-950) and pET24b(+)-ORP2(49-480) plasmids were purchased from Genscript. The pET22b_His6_STARD1(66-284) and pET22b_His6_STARD3(216-444) plasmids were a gift from James H. Hurley (University of California).^9^ The pHIS_2His6_Thrombin_STARD4(2-205;C75S)^25^ plasmid was a gift from Young Jun Im (Chonnam National University). STARD5A was a gift from Nicola Burgess-Brown (Addgene plasmid #42392; http://n2t.net/addgene:42392; RRID:Addgene_42392).

### Protein expression and purification

The ASTER domains of human Aster-A(359-547), -B(364-552) and -C(318-504) in pGEX-6p-2rps vectors including an N-terminal PreScission-cleavable GST-tag were expressed in *Escherichia coli* (*E. coli*) OverExpress C41 in Terrific Broth (TB) medium for 16 h at 18 °C after the induction with 0.1 mM IPTG. Cells were harvested at 3,500g for 15 min and lysed by sonication in buffer containing 50 mM HEPES pH 7.5, 300 mM NaCl, 10% (v/v) glycerol, 5 mM DTT, 0.1% (v/v) Triton X-100 and protease inhibitor mix HP plus (Serva). The lysate was purified by affinity chromatography on a GSTrap FF column (Cytiva) using an ÄKTA Start (Cytiva) in buffer containing 50 mM HEPES pH 7.5, 300 mM NaCl, 10% (v/v) glycerol, 5 mM DTT and 0.01% (v/v) Triton X-100. The GST-tag was cleaved overnight on the column at 4 °C. The Aster sterol binding domains were further purified by size-exclusion chromatography (SEC) on a HiLoad 16/600 Superdex 75 pg (Cytiva) in buffer containing 20 mM HEPES pH 7.5, 300 mM NaCl, 10% (v/v) glycerol and 2 mM DTT.

The START domains of human STARD1(66-284), STARD3(216-444), STARD4(2-205; C75S) and STARD5(6-213) harboring an N-terminal His_6_-Tag were expressed in *E. coli* BL21(DE3) in Luria-Bertani Broth (LB) medium for approximately 16 h at 18 °C after induction with 0.15 mM IPTG. Cells were harvested at 3,500g for 15 min and lysed by sonication in buffer containing 50 mM HEPES pH 7.5, 150 mM NaCl, 5% (v/v) glycerol, 5 mM DTT, 0.1% (v/v) Triton X-100 and EDTA-free protease inhibitor cocktail (Sigma-Aldrich). The cleared lysate was purified by affinity chromatography on a Ni-NTA Superflow Cartridge (Qiagen) using an ÄKTA Start (Cytiva) in buffer containing 50 mM HEPES pH 7.5, 150 mM NaCl, 5 % (v/v) glycerol, 5 mM DTT. START domains were eluted by using elution buffer containing 50 mM HEPES pH 7.5, 150 mM NaCl, 5% (v/v) glycerol, 5 mM DTT and 500 mM imidazole. Proteins were further purified by SEC on a HiLoad 16/600 Superdex 75 pg (Cytiva) in buffer containing 20 mM HEPES pH 7.5, 150 mM NaCl, 5% (v/v) glycerol and 2 mM DTT.

The ORP domain of human OSBP(377-807) in the pGEX-6p-1 vector with an N-terminal PreScission-cleavable GST-tag was expressed in *E. coli* OverExpress C41 in LB medium for 16 h at 18 °C after induction with 0.1 mM IPTG. Cells were harvested at 3,500g for 15 min and lysed by sonication in buffer containing 20 mM HEPES pH 7.5, 300 mM NaCl, 10% (v/v) glycerol, 5 mM DTT, 0.1% (v/v) Triton X-100 and EDTA-free protease inhibitor cocktail (Sigma-Aldrich). The cleared lysate was purified by affinity chromatography on a GSTrap HF column (Cytiva) using an ÄKTA Start (Cytiva) in buffer containing 20 mM HEPES pH 7.5, 300 mM NaCl, 10% (v/v) glycerol, 5 mM DTT. OSBP(377-807) was eluted by using elution buffer containing 20 mM HEPES pH 7.5, 300 mM NaCl, 10% (v/v) glycerol, 5 mM DTT and 10 mM reduced glutathione. Proteins were further purified by SEC on a HiLoad 16/600 Superdex 75 pg (Cytiva) using an ÄKTA Explorer (Cytiva) in buffer containing 20 mM HEPES pH 7.5, 150 mM NaCl, 10% (v/v) glycerol and 2 mM DTT.

The ORP domains of human ORP1(534-950) in pET24b(+) and ORP2(49-480) in the pET24b(+) vector including an N-terminal His6-Tag were expressed in *E. coli* BL21(DE3) in TB medium for approximately 16 h at 18 °C after induction with 0.1 mM IPTG. Cells were collected at 3,500 g for 15 min and lysed by sonification in buffer containing 10 mM Tris-HCl pH 8, 300 mM NaCl, 5% (v/v) glycerol, 2 mM DTT, 0.1% (v/v) Triton X-100 and EDTA-free protease inhibitor cocktail (Sigma-Aldrich). The cleared lysate was purified by affinity chromatography on a Ni-NTA Superflow Cartridge (Qiagen) using an ÄKTA Start (Cytiva) in buffer containing 10 mM Tris-HCl, pH 8, 300 mM NaCl, 5% (v/v) glycerol, 2 mM DTT. ORP domains were eluted using elution buffer containing 10 mM Tris-HCl, pH 8, 300 mM NaCl, 5% (v/v) glycerol, 500 mM imidazole, 2 mM DTT. The proteins were further purified by SEC on a HiLoad 16/600 Superdex 75 pg (Cytiva) using an ÄKTA Explorer (Cytiva) in buffer containing 10 mM Tris-HCl, pH 8, 150 mM NaCl, 5% (v/v) glycerol and 2 mM DTT.

### UV-Vis Absorbance and Emission Spectroscopy

Measurements were performed on a Tecan Spark Cyto spectrophotometer in an integrated JGS2 quartz 96-well plate from MicQuartz. The measurements were carried out in three different solvents: DCM, methanol and buffer (20 mM HEPES pH 7.5, 300 mM NaCl, 2 mM DTT), which were used at a final concentration of 40-60 µM, in 200 µL samples. Absorption and emission spectra were corrected to solvent blanks. Absorption measurements are corrected for pathlength, and emission spectra are then normalized to account for variations in gain. All measurements were performed in duplicate and at 25 °C.

### Fluorescence polarization

Fluorescence emission measurements as well as fluorescence intensity and polarization experiments were performed in a buffer composed of 20 mM HEPES pH 7.5, 300 mM NaCl, 0.01% (v/v) Tween-20, 0.5% (v/v) glycerol and 2 mM DTT in a final volume of 30 µl in black, flat-bottom, non-binding 384-well plates (Corning). Excitation and emission for each sterol fluorophore are available in Supplementary Table 2.

Fluorescence polarization experiments were performed in a buffer containing 20 mM HEPES pH 7.5, 300 mM NaCl, 0.01% (v/v) Tween-20, 0.5% glycerol and 2 mM DTT in a final volume of 30 µl in black, flat-bottom, non-binding 384-well plates (Corning). For k_d_ measurements fluorophore was incubated with desired concentrations of protein. For competition experiments, 20 nM 22-NBD-cholesterol or 80 nM 25-NBD-cholesterol was mixed with protein and incubated with desired concentrations of screening compounds. The fluorescence polarization signal was measured using a Spark Cyto multimode microplate reader (Tecan) with filters set at 485 ± 20 nm for excitation and at 535 ± 20 nm for emission. The data was analyzed using GraphPad Prism 5. Measured mP values were normalized setting 100% inhibition as the FP signal from the protein + fluorophore control well and 0% as the FP signal from the fluorophore control well. Curves were fitted to the normalized data via non-linear regression to allow the determination of IC_50_ values. The assay conditions for each protein are available in Supplementary Table 3.

### Fluorescence Intensity (FI) assay

#### Fluorophore Titrations

FI experiments were performed in a buffer containing 20 mM HEPES pH 7.5, 300 mM NaCl, 0.01% (v/v) Tween-20, and 2 mM DTT in a final volume of 30 µl in black, flat-bottom, non-binding 384-well plates (Corning). For k_d_ titrations of protein against fluorophore: fluorophore concentration was kept constant at either 100 nM or 200 nM. The protein was then diluted in a 3-fold fashion from 15 µM or 5 µM. 15 µL of both the fluorophore solution and the protein solution were added to each well and then allowed to incubate for 20 minutes after centrifuging. A protein only control titration is made as a control that will be normalised against. The fluorescence intensity signal was measured using a Spark Cyto multimode microplate reader (Tecan) with monochromator set at 280 ± 5 nm and 324 ± 5 nm for excitation and at 450 ± 15 nm for emission. The data was analysed using GraphPad Prism 5. Measured FI values were normalized against the protein only control titration. Curves were fitted to the normalized data via non-linear regression to allow the determination of k_d_ values.

#### STARD4 competitive assay

In the competitive setup for STARD4, competitor ligands were transferred to the Corning 384-well plate using the LabCyte Echo 550 Liquid Handler. In a single concentration screen the final ligand concentration was 10 µM and in dose response a 2-fold dilution across 8 points was used starting with 20 or 10 µM. The final concentration of fluorophore and protein was 80 nM and 120 nM, respectively, with the 27-HCTL and STARD4 stocks being pre-incubated on ice, before adding 30 µL to the plate containing competitor ligands. The plate was then centrifuged and incubated for 20 minutes, at room temperature, before reading. The fluorescence intensity signal was measured using a Spark Cyto multimode microplate reader (Tecan) with monochromator set at 280 ± 5 nm and 324 ± 5 nm for excitation and at 450 ± 15 nm for emission. The data was analysed using GraphPad Prism 5. Measured FI values were normalized setting 100% inhibition as the FI signal from the protein only control well and 0% as the FI signal from the protein + fluorophore control well. Curves were fitted to the normalized data via non-linear regression to allow the determination of IC_50_ values.

### Differential Scanning Fluorimetry

Differential scanning fluorimetry (DSF) experiments were performed in a buffer composed of 20 mM HEPES pH 7.5, 300 mM NaCl, and 2 mM DTT in Milli-Q water. Stock solutions of STARD3 and STARD5 were made at a concentration of 5 µM, and STARD4 at 2.5 µM in the HEPES Buffer. A LabCyte Echo 550 Liquid Handler was used to transfer the required amount of DMSO dissolved ligand into the 384-well plate (LightCycler® 480 Multiwell Plate 384, white). Final concentrations of ligands in a single concentration high-throughput screening are 12.5 µM. For dose response, a 2-fold dilution over 8 points was made starting at concentrations of either 100 µM (STARD3 and 5) or 50 µM (STARD4). This was lowered for compounds clearly showing solubility issues and became compound specific. After compound addition, 10 µL of protein solutions was manually pipetted to each well using an electronic 12-channel pipette. The plate was then briefly centrifuged before subsequently adding 20 nL of 5000x SYPRO orange (Sigma-Aldrich), with the Echo liquid handler, for a final concentration of 10x SYPRO orange. The fluorescence intensity was measured in a Roche LightCycler 480 II with an initial incubation at room temperature for 10 minutes before ramping the temperature from 30 °C, by steps of 0.2 °C, up to 90 °C with incubation for 5 seconds at each step. Melting temperatures were calculated with the Roche TSA analysis program.

### Circular Dichroism (CD) assay

All proteins were buffer exchanged to a PBS buffer (10 mM Phosphate-buffered saline (PBS), 138 mM NaCl, 2.7 mM KCl, pH 7.4 in Milli-Q water). The protein concentrations measured are 2.5 µM. In the cases of ligand-protein measurements, protein concentration was maintained at 2.5 µM in the presence of 5 µM ligand. The instrument used for the analysis is a Jasco J-1500 CD Spectrometer. Samples of 200 µL were measured in Quartz SUPRASIL 1 mm Cuvettes (Hellma Analytics).

#### CD Spectra

For the CD Spectra, measurements of CD (mdeg), HT (V), and Absorbance (A.U.) were recorded from 250 – 190 nm, every 1 nm. All measurements were done in duplicate per sample resulting in an averaged measurement. Cell temperature was maintained at 25 °C, with a D.I.T. of 2 seconds, bandwidth of 1 nm and scanning speed of 50 nm/min. CD Spectra were not baseline corrected and instead reported alongside the PBS blank using Prism 5 (SI Figure 2 and 4).

#### CD Variable Temperature Measurements (CD-VTMs)

For CD-VTMs, measurements of CD (mdeg), HT (V), LD (dOD), and Absorbance (A.U.) were recorded at the minima wavelength for each protein (e.g. 210 nm for ORP1), for a range of temperatures from 30 – 75/85 °C. The temperature was increased at a rate of 1 °C/min with a D.I.T. of 2 seconds and bandwidth of 1 nm. The melting temperatures were obtained using the Jasco instrument analysis software whilst the spectra were replotted using Prism 5 to illustrate the unfolding of the secondary structure (SI Figure 4).

### Sterol transfer assay

#### Preparation of Vesicles

##### TopFluor Setup

1,2-dioleoyl-sn-glycero-3-phosphocholine (DOPC, Avanti Polar Lipids, 850375C) was prepared in chloroform (10 mg/mL); 23-(dipyrrometheneboron difluoride)-24-norcholesterol (TopFluor® Cholesterol, Avanti Polar Lipids, 810255) and N-(lissamine rhodamine B sulfonyl)- 1,2-dihexadecanoyl-sn-glycero-3-phosphoethanolamine (triethylammonium salt) (Rh-DHPE, Invitrogen, L1392) were prepared in methanol (100 µM). The acceptor liposomes (LA) consist of DOPC only while the donor liposomes (LD) consist of a mixture of DOPC:TF-Chol:Rh-DHPE (99:0.5:0.5). The solvent was evaporated under a stream of nitrogen, followed by drying under vacuum overnight. The lipid films were hydrated to a final concentration of 60 µM using buffer containing 20 mM HEPES pH 7.5, 300 mM NaCl and 2 mM DTT. To fully dissolve the lipid films the solutions were vortexed and sonicated for 5 minutes in a 40 °C water bath, followed by five freeze and thaw cycles in liquid nitrogen. Extrusion through a polycarbonate membrane (13 times, 0.1 µM pore size, Avanti Polar Lipids) at 40 °C yielded homogenous unilamellar vesicles, which were kept on ice and used on the same day.

##### DHE Setup

A 2 mM stock solution of DHE in absolute ethanol was prepared from 1 mg solid (Avanti Polar Lipids, #810253). Dansyl-PE (1 mL, 1 mg/mL) in chloroform was obtained from Avanti Polar Lipids (#810333A). A stock solution of DOPC in chloroform (10 mg/mL) had previously been prepared from a 25 mg/mL solution (Avanti Polar Lipids #850375C). The following steps all took place in glass-vials covered in aluminium foil, to keep the light sensitive lipids in the dark as much as possible: The stock solutions were mixed in a molar ratio of 90/10 DOPC/DHE for the donor vesicles and 97.5/2.5 Dansyl-PE for the acceptor vesicles to a final volume of 1 mL in chloroform. Evaporation of the solvent under a stream of nitrogen, followed by drying under vacuum overnight afforded the dried lipid films. The lipid films were hydrated in a buffer of 20 mM HEPES pH 7.5, 300 mM NaCl, and 2 mM DTT to a final concentration of 260 µM. The solutions of the lipid films were vortexed extensively until full hydration was observed and sonicated for five min in a 40 °C water bath followed by five freeze-thaw cycles (-196 °C è 40 °C). Homogeneous unilamellar vesicles were obtained by extrusion 13 times through a polycarbonate membrane (0.1 µm pore size, Avanti Polar Lipids) at 40 °C. Solutions were kept on ice and used on the same day as preparation.

#### Microplate-based cholesterol transfer assay

In a non-binding clear-bottom 96-well plate (Greiner Bio-one, cat# 655906) wells were prepared as follows:

##### Topfluor Setup

**Preparation for run**: A master mix of donor and acceptor liposomes was made in buffer (20 mM HEPES pH 7.5, 300 mM NaCl, 2 mM DTT) affording a final assay concentration for both donor and acceptor as 8 µM (total liposome concentration 16 µM) and a final assay volume of 100 µL. For compound containing runs, protein and compound were incubated at a concentration 20 times the desired assay concentration for 15 min.

**The run**: 95 µL of liposome mixture was then added to each well (maximally 6 wells per run). Fluorescence intensity measurements were performed in a Tecan Spark Cyto plate reader at 25 °C, measuring from the bottom at 10 sec intervals. The excitation filter was set at 485 ± 25 nm and the emission filter was set to 590 ± 20 nm. After approximately 2 minutes, the measurement was paused, the plate was ejected, and 5 µL protein (or protein + compound pre-mix) was added as quickly as possible and mixed with a pipette to a desired final concentration. The measurement was continued, and the total measuring time was 12 min. Data was normalised to I_0_ of the donor + acceptor (before adding protein) and plotted in GraphPad Prism 5.

##### DHE Setup

**Preparation for run**: A master mix of donor and acceptor liposomes was made in buffer (20 mM HEPES pH 7.5, 300 mM NaCl, 2 mM DTT) affording a final assay concentration for both donor and acceptor as 12.5 µM (total liposome concentration 25 µM) and a final assay volume of 100 µL. For compound containing runs, protein and compound were incubated at a concentration 20 times the desired assay concentration for 15 min.

**The run**: 95 µL of liposome mixture was then added to each well (maximally 6 wells per run). Fluorescence intensity measurements were performed in a Tecan Spark Cyto plate reader at 25 °C, measuring from the bottom at 10 sec intervals. The excitation filter was set at 340 ± 20 nm and the emission filter was set to 535 ± 20 nm. After approximately 2 minutes, the measurement was paused, the plate was ejected, and 5 µL protein (or protein + compound pre-mix) was added as quickly as possible and mixed with a pipette to a desired final concentration. The measurement was continued, and the total measuring time was 12 min. Data was normalised to I_0_ of the donor + acceptor (before adding protein) and plotted in GraphPad Prism 5. The final assay setup and final assay concentrations for each protein are available in Supplementary Table 3.

#### Molecular Modelling

Software: Maestro version 13.3.121, MMshare Version 5.9.121, Release 2022-3, Platform Windows-x64. Crystal structure PDB source files were obtained from the Protein Data Bank (PDB IDs in SI Table 5). Protein preparation was carried out using the default workflow with a few minor adjustments. Namely, in the Preprocess workflow, create disulphide bonds, fill in missing loops (using Prime) were selected, and setting the variation of het states (using Epik) to pH range 7.5 ± 0.5. For the H-bond assignments workflow; H-bonds were assigned using PROPKA at pH 7.5. In the final Minimize and Delete waters workflow; a restrained minimization was performed with a convergence of 0.3 Å to heavy atoms. If there existed waters in the binding site, the preparation was optimized by running a validation test on the binding of the cognate ligands. Ligand preparation was carried out on all possible stereoisomeric forms of the ligand, which is desalted, and ionized at pH 7.5 ± 0.5 using Epik. Force field used is OPLS4.

For initial screenings of crystal structures with ligands, receptor grid generation was carried out centroid on the co-crystalised ligand. High throughput ligand docking was carried out using Glide. Utilising standard precision (SP), with flexible ligands. The settings allowed for sampling of nitrogen inversions and ring conformations. Epik state penalties were applied to docking scores. Three poses for each ligand were generated, allowing for more precise accounts of ligand-protein viability. Post-docking minimization and strain correction were also applied to the scoring of each ligand. Pose views were sampled in correlation to the Docking scores. For Induced Fit workflows, the binding domain is centroid on the resident ligand. Ligands are free to sample variations in ring conformation. In glide docking the protein preparation constrained refinement is selected for with maximally 20 poses to be generated. Prime refinement is within 5 Å of ligand poses and the Glide redocking is at standard precision. Pose analysis was performed on all examples provided from the simulation. Key considerations were on the retention of any significant position, residue interactions and orientations that were observed as median averages. Details for the used crystal structures of each protein are available in Supplementary Table 5.

## Notes

### Competing Interest Statement

The authors have declared no competing interest.

