## Supporting Information for "Endogenous and fluorescent sterols reveal the molecular basis for ligand selectivity of human sterol transporters"

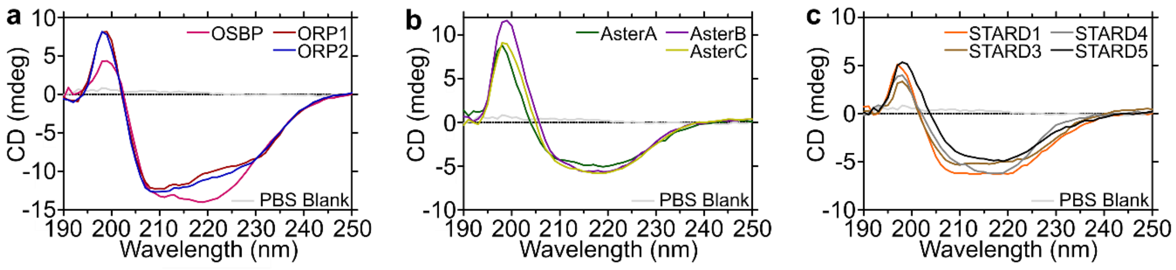


**Supplementary Figure 1**: CD spectra of 2.5 µM sterol transport proteins in PBS Buffer (pH 7.5); **a** – OSBP, ORP1, and ORP2; **b** – Aster-A, Aster-B, and Aster-C; **c** – STARD1, STARD3, STARD4, and STARD5, illustrating the correct folding of the proteins' secondary structures. One representative experiment from three replicates shown.

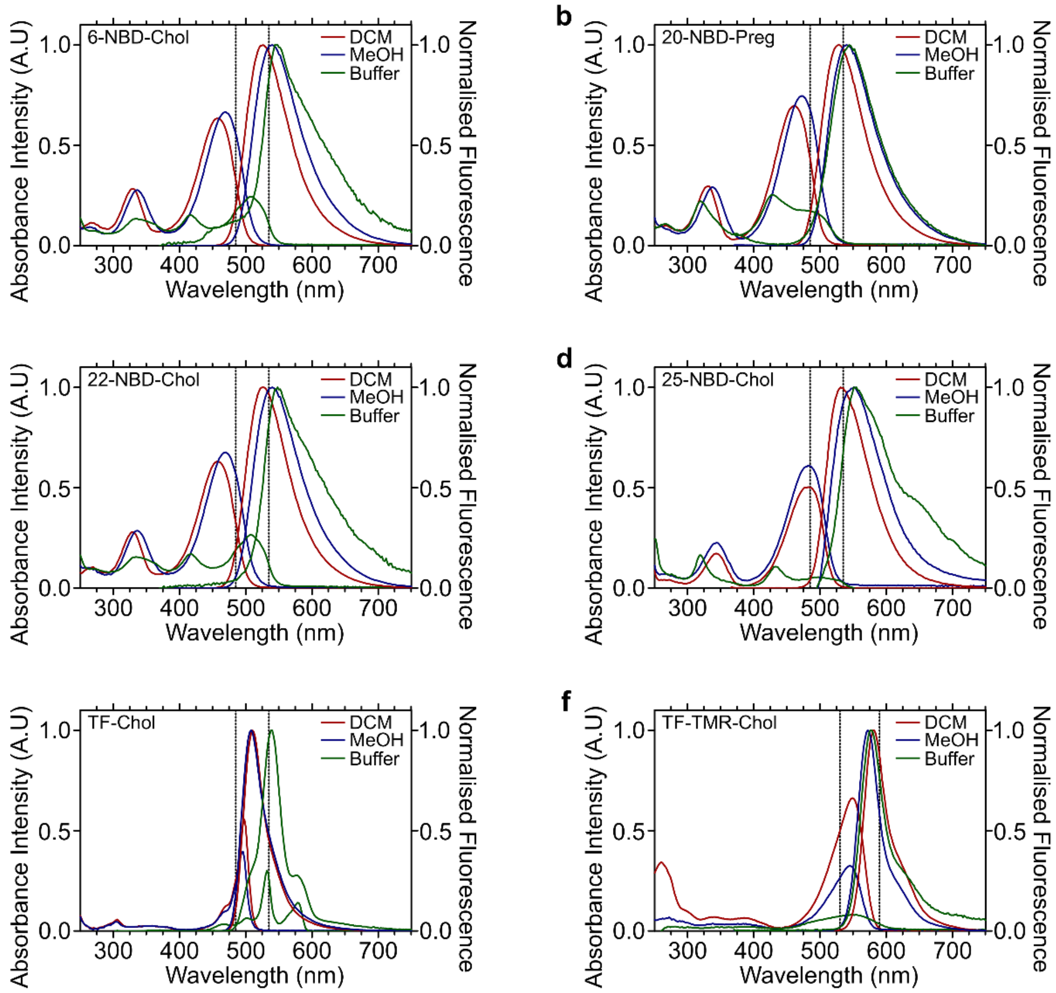


**Supplementary Figure 2**: Absorbance and emission spectra in DCM, MeOH or buffer (20 mM HEPES, pH 7.5, 300 mM NaCl) for **a** – 6-NBD-Chol, **b** – 20-NBD-Preg, **c** – 22-NBD-Chol, **d** – 25-NBD-Chol, **e** – TF-Chol, and **f** – TF-TMR-chol. One representative experiment from three replicates shown.

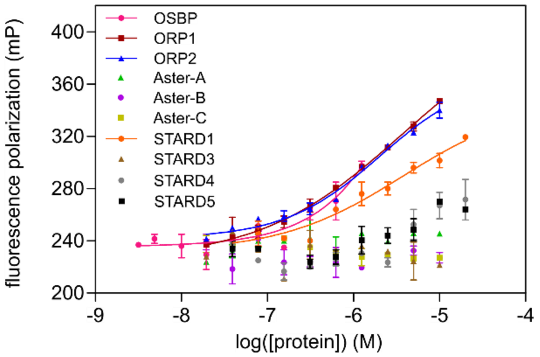


**Supplementary Figure 3:** Direct titration of 6-NBD-chol against increasing protein concentrations enables the determination of $k_d$ by monitoring changes in fluorescence polarization. Experimental points were measured in duplicates on each plate and were replicated in n = 3 biologically independent experiments. Error bars indicate s.e.m.

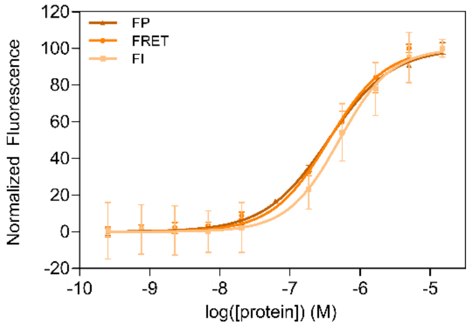


**Supplementary Figure 4:** Direct titration of 20-NBD-preg against increasing STARD1 concentrations as assessed by FI, FP and FRET. For a direct comparison the data are normalized to the highest and the lowest value in each measurement. Experimental points were measured in duplicates on each plate and were replicated in n = 3 biologically independent experiments. Error bars indicate s.e.m..

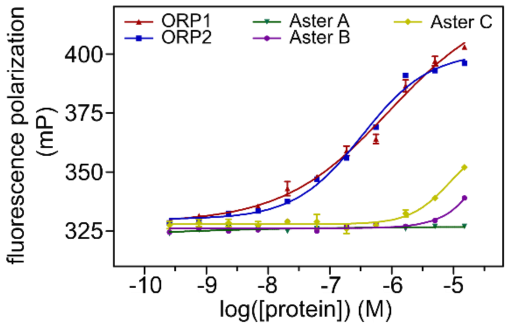


**Supplementary Figure 5:** Differences in fluorescence polarization upon titration of SYPRO Orange against increasing protein concentrations showing binding of ORP1 and ORP2 to SYPRO Orange. Experimental points were measured in duplicates on each plate and were replicated in n = 3 biologically independent experiments. Error bars indicate s.e.m..

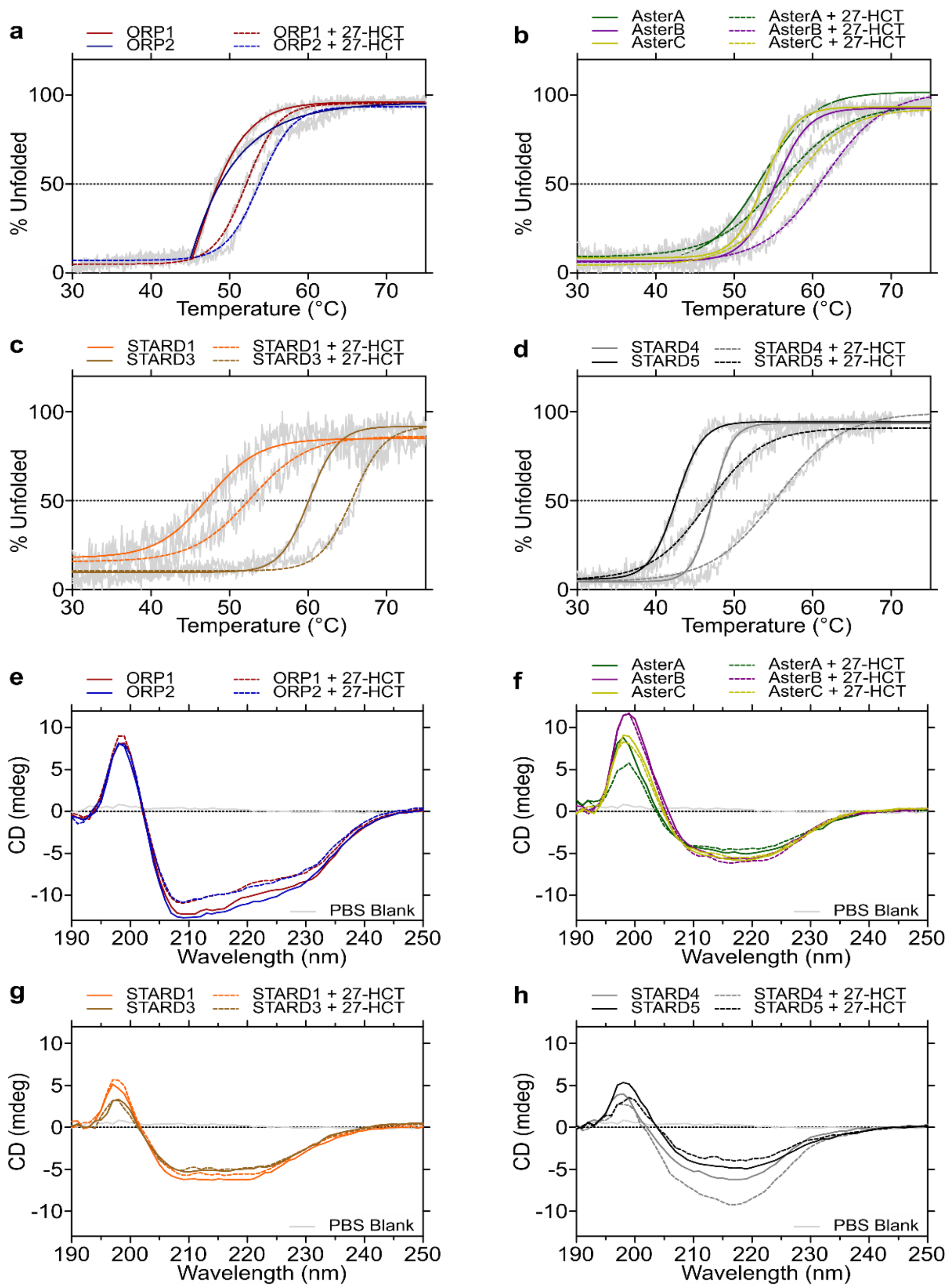


**Supplementary Figure 6**: CD Variable Temperature Measurements & CD spectra of 2.5 µM sterol transport proteins in the absence (solid lines) and presence of 5 µM 27-HCT (dashed lines) in PBS Buffer (pH 7.5); **a & e** – ORP1 and ORP2; **b & f** – Aster-A, Aster-B, and Aster-C; **c & g** – STARD1 and STARD3; **d & h** - STARD4 and STARD5.

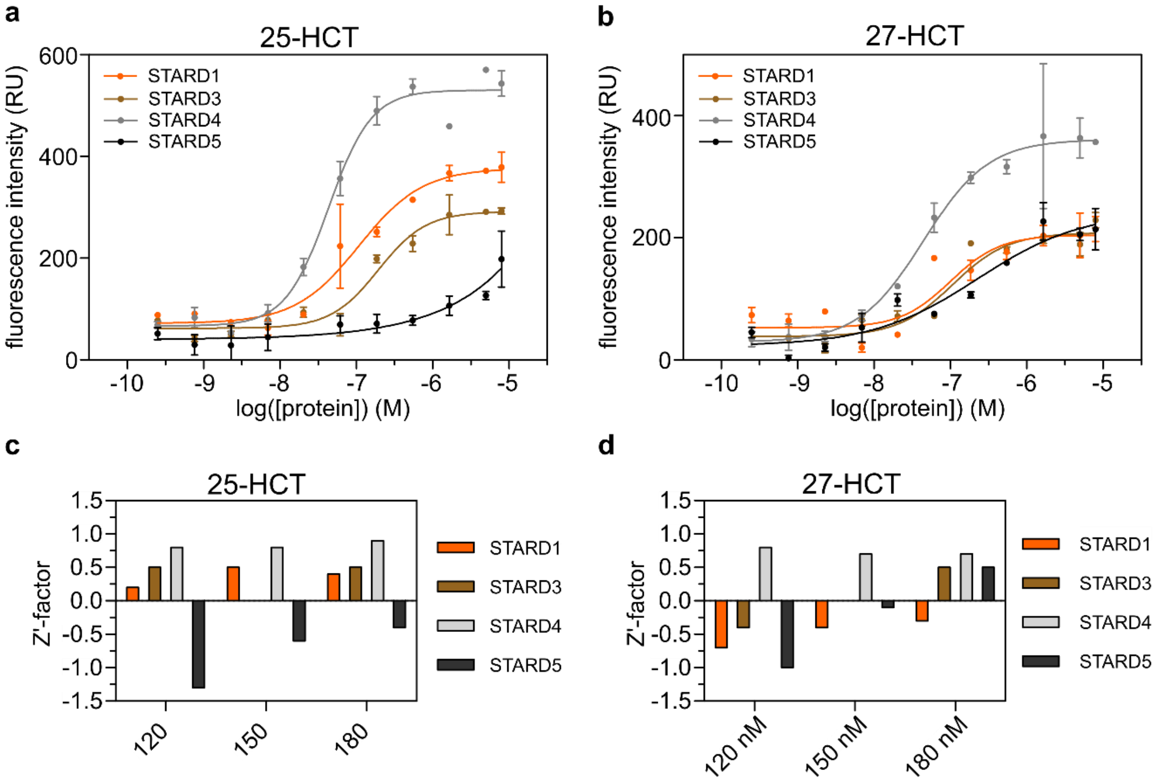


**Supplementary Figure 7:** Fluorescence intensity titrations of 25-HCTL (**a**) and 27-HCTL (**b**) against the STARD-domain containing proteins. Excitation in these measurements was carried out at 324 ± 10 nm and emission measured at 450 ± 10 nm. Panels **c** (25-HCTL) and **d** (27-HCTL) represent the Z'-factor calculations of assay stability and performance. Z'-factor calculations were measured on assay conditions of 80 nM HCTL fluorophore concentration and 120, 150, and 180 nM protein concentrations.

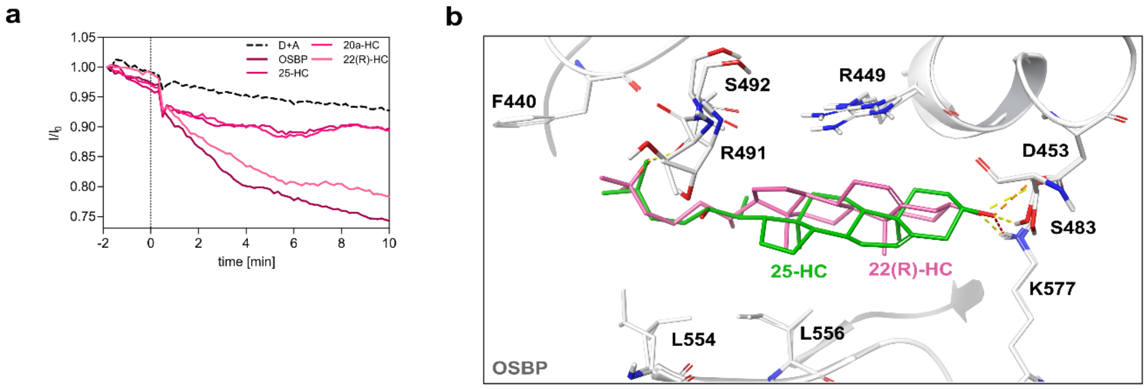


**Supplementary Figure 8:** Oxysterols bind and inhibit OSBP. **a** Inhibition of sterol transport mediated by OSBP (500 nM) by different oxysterols. One representative experiment is shown from two independent experiments (n=2), D = donor liposomes, A = acceptor liposomes. **b** Comparison of the predicted binding poses of 25-HC and 22(*R*)-HC into the recently published crystal structure of OSBP (pdb: 7v62) highlighting crucial interactions of 25-HC with the OSBP-ORD as well as possible clashes induced by the 22-OH in (*R*)-configuration.

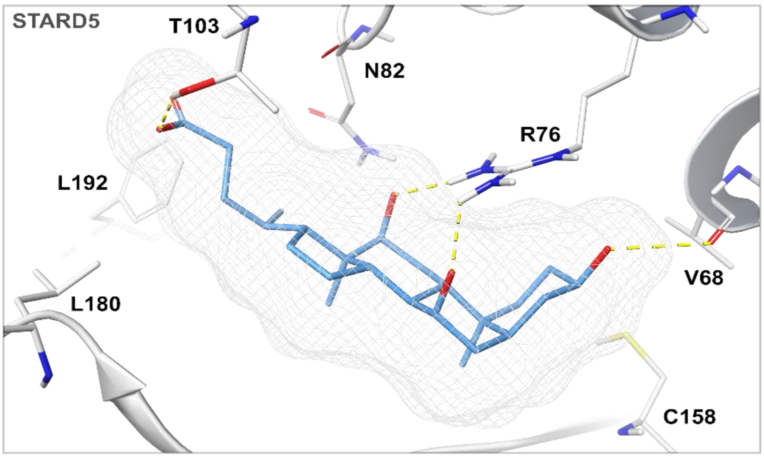


**Supplementary Figure 9:** Representation of the predicted binding pose of CA into the published crystal structure of STARD5 (pdb: 2r55) highlighting crucial interactions of CA with R76 and V68.

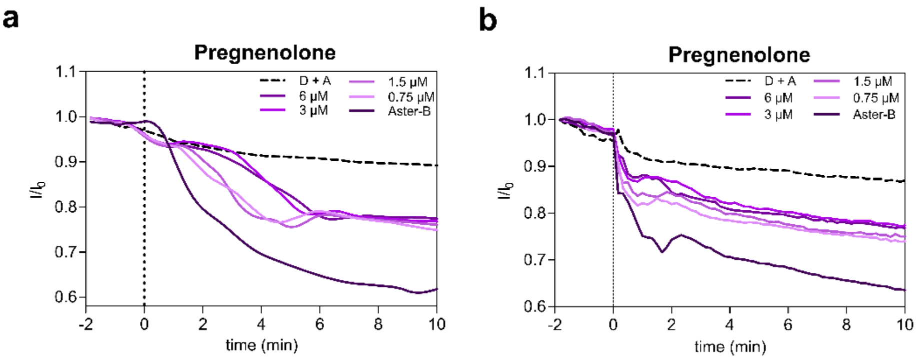


**Supplementary Figure 10: a** Inhibition of sterol transport mediated by 125 nM Aster-B by different concentrations of pregnenolone. One representative experiment is shown from two independent experiments (n=2), D = donor liposomes, A = acceptor liposomes. The compound was incubated with the liposomes before the protein was added after 2 min. **b** Inhibition of sterol transport mediated by 125 nM Aster-B by different concentrations of pregnenolone. One representative experiment is shown from two independent experiments (n=2), D = donor liposomes, A = acceptor liposomes. The compound was incubated with Aster-B before they were added after 2 min.

**Supplementary Table 1: Summary of the in this study tested natural products, supplier, CAS number and catalogue number.**

| Compound name | Abbreviation | Company | CAS Number | Catalogue Number |
|---|---|---|---|---|
| **6-NBD-cholesterol** | 6-NBD-chol | Avanti Lipids | **201731-19-3** | 810251P |
| **20-NBD-pregnenolone** | 20-NBD-preg | | Synthesised in-house | |
| **22-NBD-cholesterol** | 22-NBD-chol | Invitrogen™ | **-** | N1148 |
| **25-NBD-cholesterol** | 25-NBD-chol | Avanti Lipids | **105539-27-3** | 810250P |
| **TopFluor-TMR-cholesterol** | TF-TMR-chol | Avanti Lipids | **2342615-73-8** | 810385P |
| **TopFluor-cholesterol** | TF-chol | Avanti Lipids | **878557-19-8** | 810255P |
| **Dehydroergosterol** | DHE | Avanti Lipids | **516-85-8** | 810253P |
| **25-hydroxycholestatrienol** | 25-HCTL | | Synthesised in-house[1] | |
| **27-hydroxycholestatrienol** | 27-HCTL | | Synthesised in-house[2] | |
| **4β-hydroxycholesterol** | 4β-HC | Synthesised in-house[3] | **17320-10-4** | |
| **7α-hydroxycholesterol** | 7a-HC | Avanti Lipids | **566-26-7** | 700034P |
| **7β-hydroxycholesterol** | 7b-HC | Avanti Lipids | **566-27-8** | 700035P |
| **7-ketocholesterol** | 7-KC | Avanti Lipids | **566-28-9** | 700015P |
| **20α-hydroxycholesterol** | 20a-HC | Avanti Lipids | **516-72-3** | 700156P |
| **22(*R*)-hydroxycholesterol** | 22(R)-HC | Avanti Lipids | **17954-98-2** | 700058P |
| **24(*S*)-hydroxycholesterol** | 24(S)-HC | Avanti Lipids | **474-73-7** | 700061P |
| **25-hydroxycholesterol** | 25-HC | Avanti Lipids | **2140-46-7** | 700019P |
| **27-hydroxycholesterol** | 27-HC | Avanti Lipids | **20380-11-4** | 700021P |
| **Cholestane-3,5,6-triol** | CT | Synthesised in-house[3] | **277329-65-4** | |
| **Estradiol** | | Sigma-Aldrich | **50-28-2** | E2758 |
| **Androstenediol** | AD | DTU compound collection | **521-17-5** | - |
| **Dehydroepiandrosterone** | DHEA | Fluorochem | **53-43-0** | 449222 |
| **Pregnenolone** | Pregn | Combi-blocks | **145-13-1** | QA-1884 |
| **21-acetoxy pregnenolone** | 21-AcP | DTU compound collection | **566-78-9** | sc-231272 |
| **Pregnenolone acetate** | | DTU compound collection | **1778-02-5** | - |
| **Progesterone 3-acetyl enol ether** | 3-AcP | DTU compound collection | **4954-06-7** | - |
| **3β,17α-dihydroxypregn-5-en-20-one 3-acetate** | | DTU compound collection | **1863-39-4** | - |
| **Dehydroepiandrosterone acetate** | DHEA ac | DTU compound collection | **853-23-6** | - |
| **Ergosterol acetate** | | DTU compound collection | **2418-45-3** | - |
| ***ent*-U18886A** | | | Synthesised in-house[4] | |
| **U18666A** | | MedChemExpress | **3039-71-2** | HY-107433 |
| **Testosterone** | Testost | TCI | **58-22-0** | T0027 |
| **17α-Methyl-6,7-dehydrotestosterone** | 17α-Me-6,7-DHT | DTU compound collection | **5585-85-3** | - |
| **4β,5β-Epoxytestosterone** | | DTU compound collection | **-** | - |
| **Progesterone** | Progest | DTU compound collection | **57-83-0** | - |
| **17α-hydroxyprogesterone** | | DTU compound collection | **68-96-2** | - |
| **pregnan-3,20-dione 4a,5a-oxide** | | DTU compound collection | **-** | - |
| **Testosterone propionate** | Testost prop | DTU compound collection | **57-85-2** | - |
| **6-Dehydrotestosterone acetate** | 6-DHT-Ac | DTU compound collection | **2352-19-4** | - |
| **4,5-epoxyandrostane-3,17-dione** | | DTU compound collection | **77057-73-9** | - |
| **Adrenosterone** | | DTU compound collection | **382-45-6** | - |
| **2,17β-diacetoxy-androst-1,4-dien-3-one** | | DTU compound collection | **-** | - |
| **Hydrocortisone** | | DTU compound collection | **50-23-7** | - |

| | | | | |
|---|---|---|---|---|
| **Cortisone 21-acetate** | | DTU compound collection | **50-04-4** | - |
| **4-cholesten-3-one** | | Avanti Lipids | **601-57-0** | 700065P |
| **Cholic acid** | CA | Avanti Lipids | **81-25-4** | 700212P |
| **Dehydrocholic acid** | DHCA | Avanti Lipids | **81-23-2** | 700215P |
| **Chenodeoxycholic acid** | CDCA | Apollo Scientific | **474-25-9** | BIB6026 |
| **Deoxycholic acid** | DCA | Avanti Lipids | **83-44-3** | 700197P |
| **Hyodeoxycholic acid** | HDCA | Combi-blocks | **83-49-8** | QA-8913 |
| **β-sitosterol** | | Avanti Lipids | **83-46-5** | 700107P |

**Supplementary Table 2: Excitation and emission for each sterol fluorophore.**

| Ligand | Ex/Em |
|---|---|
| **NBD-cholesterol analogues** | 485(20) nm/ 535(25) nm |
| **TopFluor cholesterol** | 485(20) nm/ 535(25) nm |
| **TopFluor-TMR-cholesterol** | 535(25) nm/ 590(20) nm |

**Supplementary Table 3: Summary of the FP assay conditions for each protein.**

| Protein | Protein conc. for selectivity measurements | Protein conc. for dose response measurements | Incubation temp. | **Incubation time** (Protein/NBD cholesterol) | **Incubation time** (Protein-NBD-cholesterol/compound) |
|---|---|---|---|---|---|
| **Aster-A** | 1 µM | 0.5 µM | 4 °C | 20 min | 20 min |
| **Aster-B** | 1 µM | 1 µM | 4 °C | 20 min | 20 min |
| **Aster-C** | 1 µM | 0.5 µM | 4 °C | 20 min | 20 min |
| **STARD1** | 1 µM | 1 µM | 4 °C | 30 min | 20 min |
| **OSBP** | 1 µM | 0.25 µM | 4 °C | 60 min | 20 min |
| **ORP1** | 1 µM | 0.2 µM | 4 °C | 60 min | 20 min |
| **ORP2** | 1 µM | 0.7 µM | 4 °C | 60 min | 20 min |

**Supplementary Table 4: Summary of the used liposome setup and protein concentration used for the sterol transport assay.**

| STP | Assay Setup | Assay Conc. (µM) |
|---|---|---|
| **OSBP** | Topfluor | 0.5 |
| **Aster A** | Topfluor | 0.125 |
| **Aster B** | Topfluor | 0.125 |
| **Aster C** | Topfluor | 0.25 |
| **STARD1** | DHE | 0.5 |
| **STARD3** | DHE | 2.0 |
| **STARD4** | DHE | 0.5 |

**Supplementary Table 5: Excitation and emission for each sterol fluorophore.**

| | PDB ID | Co-crystalised ligand |
|---|---|---|
| **OSBP** | 7v62 | Cholesterol |
| **ORP1** | 5zm5 | Cholesterol |
| **ORP2** | 5zm8 | PI(4,5)P2 |
| **STARD1** | 3p0l | no ligand |
| **STARD3** | 5i9j | no ligand |
| **STARD4** | 6l1d | no ligand |
| **STARD5** | 2r55 | no ligand |
| **AsterA** | 6GQF | no ligand |
| **AsterB** | Homology Model vs Aster A | no ligand |
| **AsterC** | 7AZN | no ligand |

### Synthesis of 20-NBD-pregnenolone (20-NBD-preg)

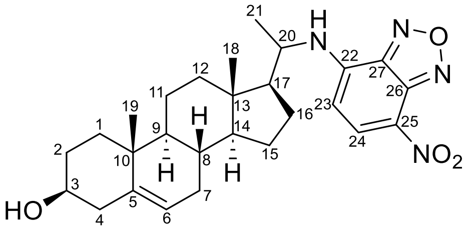

Pregnenolone (209 mg, 0.66 mmol, 1.0 mol. equiv.) and ammonium acetate (506 mg, 6.56 mmol, 10.0 mol. equiv.) were dissolved in 10 mL MeOH, to which sodium cyanoborohydride (62 mg, 8.98 mmol, 1.5 mol. equiv.) was added. The reaction mixture was stirred at room temperature for 18 hours over 4 Å mol. sieves until reaction was completed by TLC (85:15:1 $Et_2O$:MeOH:$NH_4OH$, stained with $KMnO_4$). The crude mixture was then filtered and diluted with 10% *i*PrOH in $CHCl_3$. The organic layer was subsequently washed with sat. $NaHCO_3$, $H_2O$, and brine, after which it was dried over $Na_2SO_4$. Removal of the solvent under reduced pressure gave the crude 20-aminopregn-5-en-3β-ol (275 mg) as a pale pink solid, which was used in the next step without any purification.

The crude 20-aminopregn-5-en-3β-ol (58 mg, 1.0 mol. equiv.) was dissolved in 1.5 mL of a 1:3 mixture (v/v) of MeOH and $CHCl_3$. To this, 0.2 mL sat. $NaHCO_3$ solution, and NBD-Cl (44 mg, 0.22 mmol, 1.2 mol. equiv.) in 0.5 mL 1:3 MeOH:$CHCl_3$ were added. The mixture was left to react at room temperature. After 40 hours, no more product formation was observed by TLC (5% acetone in DCM), and the reaction was stopped. The reaction mixture was diluted with 5% *i*PrOH in $CHCl_3$, and subsequently washed with 0.5 M HCl and brine. The organic layer was then dried over $Na_2SO_4$, after which the solvent was removed in vacuo. Using flash column chromatography, the crude product was purified. First, in a column in 1% acetone in DCM, then a second column in 20% EtOAc in n-heptane. This afforded 20-NBD-pregnenolone as a dark orange solid (22 mg, 0.047 mmol, 26% over two steps).

**TLC**: $R_f$ = 0.49 (5% acetone in DCM)

**$^1$H NMR** (400 MHz, $CDCl_3$) δ 8.50 (d, J = 8.7 Hz, 1H, H24), 6.18 (d, J = 8.7 Hz, 1H, H23), 6.13 (d, J = 9.4 Hz, 1H, NH), 5.35 – 5.33 (m, 1H, H6), 3.80 – 3.75 (m, 1H, H20), 3.51 (tt, J = 11.0, 4.5 Hz, 1H, H3), 2.32 – 2.17 (m, 2H, H4), 2.03 – 1.91 (m, 2H, H16a, H8a), 1.87 – 0.96 (m, 22H), 0.95 (s, 3H, H19) , 0.67 (s, 3H, H18).

**$^{13}$C NMR** (101 MHz, $CDCl_3$) δ 144.6 (C26), 144.2 (C27), 142.6 (C25), 141.0 (C5), 136.9 (C24), 123.6 (C22), 121.4 (C6), 97.9 (C23), 71.8 (C3), 56.6 (C17), 56.2 (C14), 52.2 (C20), 50.0 (C9), 42.6 (C4), 42.3 (C13), 39.5 (C12), 37.3 (C1), 36.6 (C10), 31.9 (C8), 31.8 (C7), 31.7 (C2), 26.8 (C15), 24.3 (C16), 21.1 (C11), 19.9 (C21), 19.5 (C19), 12.8 (C18).

**LRMS** (ESI+) *m/z*: 481.2 found for $[M+H]^+$, 481.28 calcd. for $C_{27}H_{37}N_4O_4^+$.

### NMR spectra

$^{1}$H spectrum of 20-NBD-pregnenolone recorded in $CDCl_3$.

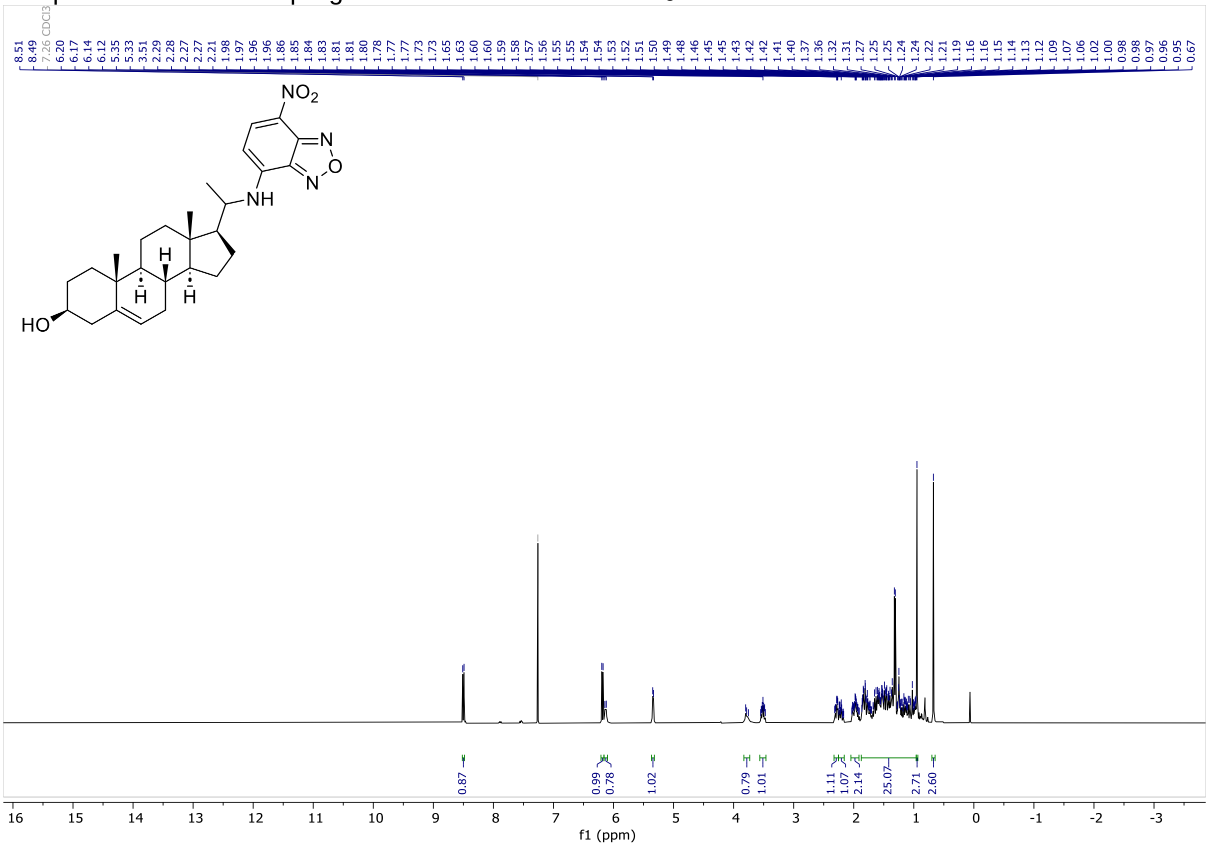


$^{13}$C spectrum of 20-NBD-pregnenolone recorded in $CDCl_3$.

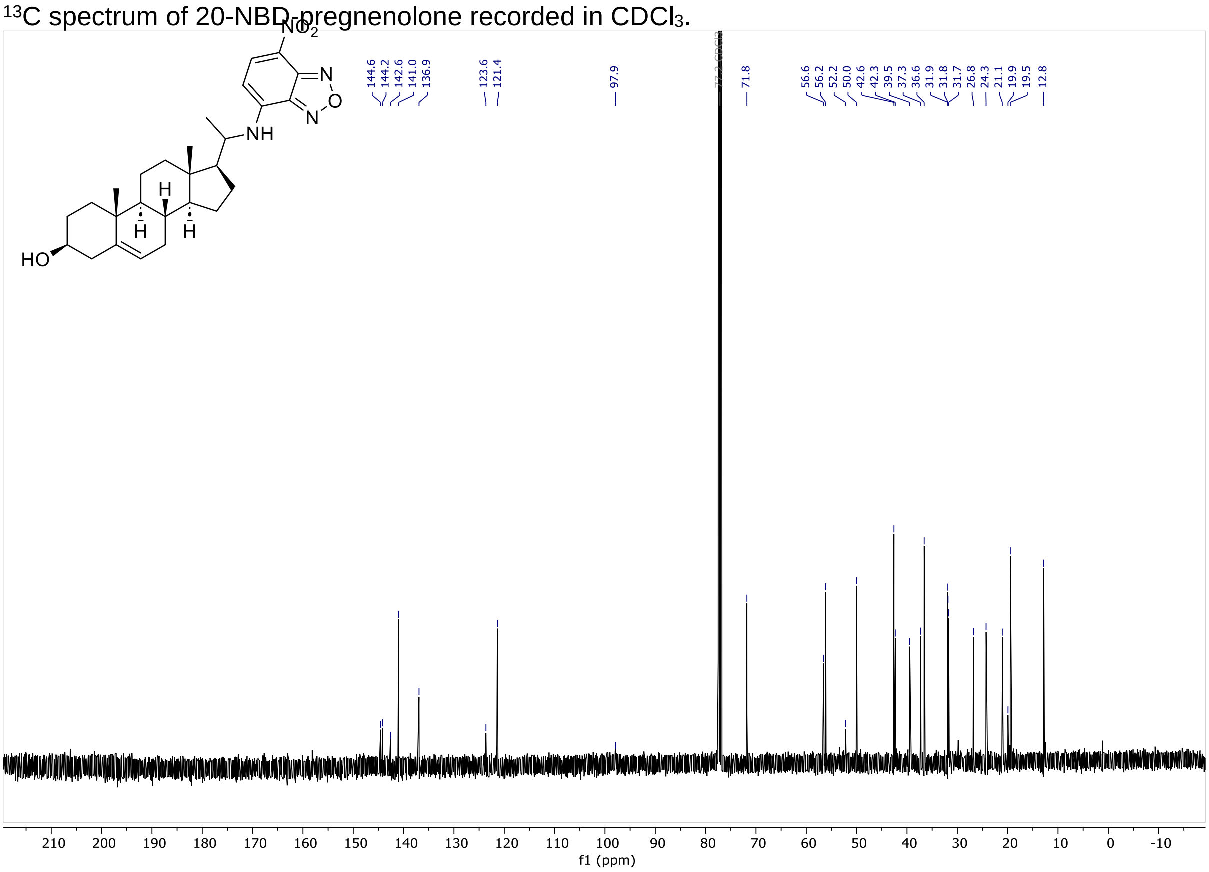
